# Population-Averaged Atlas of the Macroscale Human Structural Connectome and Its Network Topology

**DOI:** 10.1101/136473

**Authors:** Fang-Cheng Yeh, Sandip Panesar, David Fernandes, Antonio Meola, Masanori Yoshino, Juan C Fernandez-Miranda, Jean M. Vettel, Timothy Verstynen

## Abstract

A comprehensive map of the structural connectome in the human brain has been a coveted resource for understanding macroscopic brain networks. Here we report an expert-vetted, population-averaged atlas of the structural connectome derived from diffusion MRI data (N=842). This was achieved by creating a high-resolution template of diffusion patterns averaged across individual subjects and using tractography to generate 550,000 trajectories of representative white matter fascicles annotated by 80 anatomical labels. The trajectories were subsequently clustered and labeled by a team of experienced neuroanatomists in order to conform to prior neuroanatomical knowledge. A multi-level network topology was then described using whole-brain connectograms, with subdivisions of the association pathways showing small-worldness in intra-hemisphere connections, projection pathways showing hub structures at thalamus, putamen, and brainstem, and commissural pathways showing bridges connecting cerebral hemispheres to provide global efficiency. This atlas of the structural connectome provides representative organization of human brain white matter, complementary to traditional histologically-derived and voxel-based white matter atlases, allowing for better modeling and simulation of brain connectivity for future connectome studies.

## Introduction

The organization of the structural connections in the human brain determines how neural networks communicate, thereby serving as a critical constraint on brain functionality and providing potential etiology for clinical pathology (Bota et al., 2015; Sporns, 2014). Characterizing this structural organization has relied on either histological slides or neuroanatomically-validated atlases based on individual subjects (Amunts et al., 2013; Ding et al., 2016); however, a comprehensive population-averaged 3-dimensional (3D) structural connectome at the macroscale level has yet to be constructed. A population-averaged connectome is critical for demonstrating representative topological interconnectivity in the general population, a stated objective of the national investment in the Human Connectome Project (Setsompop et al., 2013; Van Essen et al., 2013). If achieved, such a map of the structural connectome could augment existing histological and single-subject atlases, thus allowing for robust modeling and simulation in both empirical and theoretical studies.

To date, diffusion MRI is the only non-invasive tool for mapping the 3D trajectories of human macroscopic white matter pathways (Fan et al., 2016; McNab et al., 2013), with preliminary success at resolving the normative pattern of several major white matter pathways (Catani et al., 2002; Guevara et al., 2012; Mori et al., 2009; Mori et al., 2008; Peng et al., 2009; Thiebaut de Schotten et al., 2011). This has been realized by resolving local fiber orientations at the voxel level and delineating entire axonal trajectories by implementing a stepwise tracking algorithm (Basser et al., 2000; Mori et al., 1999; Wedeen et al., 2012). Nonetheless, there are several caveats to the success of diffusion MRI fiber tracking, including the identification false tracts and suboptimal coverage of small pathways or those with complex geometry (Reveley et al., 2015; Thomas et al., 2014). Indeed, the validity of tractography can range from 3.75% to 92% due to differences in reconstruction methods and tracking algorithms (Maier-Hein et al., 2016). Improving the quality of resolved fiber pathways using diffusion MRI can be achieved by high-angular-resolution modalities (Glasser et al., 2016), a template averaged across a large number of subjects to facilitate fiber tracking (Yeh and Tseng, 2011), and neuroanatomical expertise to resolve errors in the automated fiber tracking process (Meola et al., 2015). Template-based approaches have been shown to reliably capture the morphological characteristics of several major white matter fascicules when validated against cadaver microdissection approaches (Fernandez-Miranda et al., 2015; Meola et al., 2016a; Meola et al., 2015; Meola et al., 2016b; Wang et al., 2016; Wang et al., 2013; Yoshino et al., 2016). Yet building a comprehensive tractography atlas of major and minor white matter pathways is still challenged by the problem of false fiber pathways, even when relying on high angular resolution data.

Here we constructed a population-averaged structural connectome, including both major and minor pathways, using an expert-vetted approach. We employed high-angular-resolution diffusion MRI data (n=842) from healthy subjects in the Human Connectome Project (HCP) database (Van Essen et al., 2012) and aggregated them into an averaged template of diffusion distributions that can inform the orientations of underlying fiber architectures. The averaged diffusion pattern of the entire sample is thus representative of non-pathological structural characteristics within healthy subjects. Based on this template, a total of 550,000 tracks were generated using a tracking method that was shown to achieve the highest number of valid connections in an open competition (Maier-Hein et al., 2016). Generated tracks were subsequently clustered and then labeled by a team of clinical neuroanatomists, capitalizing on their previous experience in both cadaveric white-matter and comparative tractography techniques (Fernandez-Miranda et al., 2015; Wang et al., 2016). Furthermore, the tracks were categorized into the projection, association, and commissural pathways to generate multi-level connectograms illustrating network topology at the macroscopic level. The strategy of this approach allowed us to compile a comprehensive atlas of the structural connectome in the human brain at the population level, allowing for taxonomical identification of pathways that together comprise the full macroscopic structural connectome.

## Methods

### Diffusion MRI acquisitions

We used the preprocessed data from Human Connectome Projects (Q1-Q4 release, 2015) acquired by Washington University in Saint Louis and University of Minnesota. A total of 842 subjects (372 males and 470 females, age 22 ~ 36) had diffusion MRI scanned on a Siemens 3T Skyra scanner using a 2D spin-echo single-shot multiband EPI sequence with a multi-band factor of 3 and monopolar gradient pulse. The spatial resolution was 1.25 mm isotropic. TR=5500 ms, TE=89.50 ms. The b-values were 1000, 2000, and 3000 s/mm^2^. The total number of diffusion sampling directions was 90, 90, and 90 for each of the shells in addition to 6 b0 images. The preprocessed data were corrected for eddy current and susceptibility artifact. The matrices for gradient nonlinearity distortion correction were used in the following diffusion MRI reconstruction.

### Super-resolution q-space diffeomorphic reconstruction

The diffusion data for each subject was reconstructed into the ICBM-152 space (ICBM: International Consortium for Brain Mapping) using the q-space diffeomorphic reconstruction (QSDR)(Yeh and Tseng, 2011), a method that conserved the diffusible spins after nonlinear transformation and could be applied to DTI, DSI, and multishell data. QSDR calculated the spin distribution function (SDF), ψ(**û**), an orientation distribution function defined as the density of spins that have diffusion displacement oriented at direction **û** during the diffusion time:

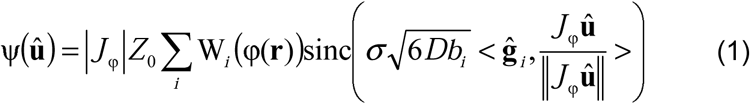

Where φ is a diffeomorphic mapping function that maps standard space coordinates **r** to the subject’s space. *J*_φ_ is the Jacobian matrix of the mapping function, whereas |*J*_φ_| is the Jacobian determinant. W_*i*_(φ(**r**)) are the diffusion signals acquired at φ(**r**). *b*_*i*_ is the b-value, and **ĝ**_*i*_ is the direction of the diffusion sensitization gradient. *σ* is the diffusion sampling ratio controlling the detection range of the diffusing spins. *D* is the diffusivity of water, and *Z*_0_ is the constant estimated by the diffusion signals of free water diffusion in the brain ventricle (Yeh and Tseng, 2011). The nonlinearity of diffusion gradients was corrected using the nonlinear terms of the magnetic field obtained from gradient coils. The HCP dataset includes a 3-by-3 gradient deviation matrix for each voxel to estimate the effective gradient direction and strength. This matrix was applied to the diffusion sensitization gradient, **ĝ**_*i*_ in Eq. (1) to correct the effect of gradient nonlinearity.

To achieve super-resolution reconstruction, we modified a nonlinear registration algorithm that used Fourier basis as the deformation function (Ashburner and Friston, 1999) to boost the registration accuracy. The original setting used a set of 7-by-9-by-7 Fourier basis at x-y-z directions for 2-mm resolution, and the computation and memory bottleneck was at the inverse of a 1327-by-1327 matrix (not a sparse matrix). We increased the resolution of the Fourier basis by 4-fold to 0.5-mm resolution (i.e. 28-by-36-by-28 Fourier basis), which required solving an 84676-by-84676 matrix for each optimization iteration. Here instead of solving the large matrix using a standard Gauss-Jordan method (a complexity of O(n^3^)), which would increase the computation time by a factor of (4x4x4)^3^=262,144, we used the Jacobi method that allowed for parallel processing and could utilize solutions from the previous iteration to speed up the processing. This greatly reduced the computation complexity to O(n) and only increased the computation time by a factor of 4x4x4=64. The parallel processing further reduced the computation time, allowing us to reconstruct the data using multi-thread resources. The final SDFs were generated at 1-mm resolution in the template space.

The registration accuracy was evaluated by the coefficient of determination (i.e., R^2^) value between each subject and template image. The distribution of the R^2^ values, as shown in Suppl. Figure 1, is skewed with a leftward tail. We therefore looked at subjects with the lowest R^2^ values at this tail for identification of outliers. This allowed us to identify two problematic datasets (#173132 and #103515) that were then reported to the HCP Consortium. It is noteworthy that we did not use the existing HCP alignment in our spatial normalization. The alignment has good point-to-point matching in the gray matter surface; however, it does not constrain or penalize large rotation of the Jacobian matrix in the white matter tissue. Consequently, the fiber architecture in the white matter can be heavily distorted to match gyral foldings. The Fourier basis used here intrinsically limits the largest possible rotation and allows for fiber tracking in the template space.

### Construction of an SDF template

Since there is no pre-existing SDF template, we first used the FMRIB 58-subject fractional anisotropy (FA) template (FMRIB, FSL) as a guiding template to normalize diffusion data of subjects. The FA map of each 842 subject was nonlinearly normalized to the FMRIB FA template to obtain the diffeomorphic mapping function. The spatial mapping functions were then used by QSDR to compute the SDFs in the standard space. The SDFs of all subjects were then averaged, voxel-by-voxel, to obtain an SDF template termed HCP-842. The computation was conducted using the cluster at Center for the Neural

Basis of Cognition, a joint Institute of Carnegie Mellon University and the University of Pittsburgh. The cluster had 24 nodes and 320 CPUs. The 842 subjects took a month of computation time to complete.

### Whole-brain tractography

We used a deterministic fiber tracking algorithm that leverages information in the SDF (Yeh et al., 2013). Each of the streamlines generated was automatically screened for its termination location. A white matter mask created by applying DSI Studio’s default anisotropy threshold to the SDF’s anisotropy values. The mask was used to eliminate streamlines with premature termination in the white matter region. We did not use ICBM T1W images as the mask because of its misalignment with FSL’s FA template.

To determine the adequate seeding density, one study showed that on average, there are around 3 fiber populations in a 2.4-mm cubic voxel (Jeurissen et al., 2013). This indicated that at least 3 seeds points are needed for each voxel with a volume of 2.4-by-2.4-by-2.4 mm^3^, which is 0.2 seeds per mm^3^. To meet the minimal requirement, we obtained 500,000 whole-brain streamlines in addition to 50,000 streamlines to cover the spinal cord connections eliminated by the white matter mask. The total number of streamlines achieved an average seeding density of 1.0 seed per mm^3^, which is 5 times of the minimum requirement.

The fiber tracking was conducted using angular thresholds of 40, 50, 60, 70, and 80 degrees. Each angular threshold generated 100,000 streamlines, and a total of 500,000 streamlines were obtained. Since the white matter mask also removed streamlines connecting to/from the spinal cord, an additional set of whole brain tracking was conducted to allow streamlines terminates at the lowest section of the brainstem. The fiber tracking was also conducted using angular thresholds of 40, 50, 60, 70, and 80 degrees. Each angular threshold generated 10,000 streamlines, and a total of 50,000 streamlines were obtained. We used different parameter combinations because different fiber trajectories are best resolved by different tractography schemes. For example, a larger angular threshold is needed for tracking fiber pathway with abrupt turning (e.g. Meyer’s loop at the optic radiation), whereas some projection pathways do not have sharp turning (e.g. corticospinal tracts) and thus can rely on lower angular thresholds. The angular threshold of 40~80 allows degrees us to capture all possible pathways.

### Initial clustering using Hausdorff distance

The tractography was clustered using single-linkage clustering. We measured the Hausdorff distance between a pair of streamlines X and Y as

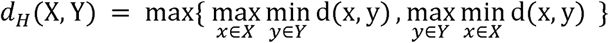

X is a set of coordinates, i.e. X={x}, whereas Y is another set of coordinates, i.e. Y={y}. d(x,y) calculates the Euclidian distance between two coordinates x and y, and the d_H_(X,Y) calculates the Hausdorff distance between set X and Y. Different Hausdorff distances were tested, and we chose 2-mm as the merging threshold to avoid over-segmentation (shorter distance) and over-merging (longer distance). The 500 largest clusters, in terms of track counts, were selected because the remaining clusters contained less than 0.01% of the total streamlines (i.e. < 50 streamlines). The same cluster selection strategy was applied to our second set of the 50,000 streamlines (i.e. the streamlines connecting to/from spinal cords), and the first 50 largest clusters were collected. Since each cluster may contain streamlines with repeated trajectories, we removed redundant trajectories that are substantially close to the one another using a Hausdorff distance of 1 mm.

### Expert labeling and examination

The 550 clusters were manually labeled by our neuroanatomy teams, including four senior neuroanatomists (JFM, AM, MY, FY) and junior neuroanatomists (DF and SP). The labeling was based on evidence from publicly available white matter atlases, existing literature, microdissection evidence, and neuroanatomy books (Table S1). The first examination round was the manual labeling conducted by 3 neuroanatomists (FY, DF, and SP). Each of the neuroanatomists independently inspected the termination locations and connecting routes of each of the 550 clusters using the 3D interface provided by DSI Studio. The anatomical label of each cluster was independently assigned and subsequently compared to identify inter-observer differences, including the naming of the cluster and whether the cluster is a false one. The inter-observer differences were found in 20 clusters (3.6% of the clusters), mostly involving the branches and segments of fiber pathways, and resolved in a joint discussion between two junior (DF, SP) and two senior neuroanatomists (FY, JFM). The clusters with the same neuroanatomy name were grouped together to form major fiber bundles. The merged bundles underwent a second round of inspection by both senior and junior neuroanatomists to identify missing branches and remove false connections. The inspection identified missing branches in anterior commissure (olfactory and occipital connections), corticothalamic tract (temporal connections), corticostriatal tract (occipital connections), corticobulbar tract, corticopontine tract (temporal and occipital connections), and tapetum of the corpus callosum. These branches were specifically tracked by a region-based approach by placing regions of interest at the target area. The final fiber bundles were subsequently categorized into the projection, association, commissural, cerebellar, brainstem, and cranial nerve pathways.

The next examination round further checked for other missing minor pathways that require a dense sampling to form a bundle. This was done by projecting the fiber bundles back to the white matter and looking for areas without track coverage. Using a region-based approach, the senior neuroanatomists (MY, AM, and FY) tracked missing minor pathways including acoustic radiation, posterior commissure, brainstem pathways such as rubrospinal tract (RST), spinothalamic tract (STT), dorsal longitudinal fasciculus (DLF), lateral lemniscus (LL), medial lemniscus (ML), and cranial nerves such as CN VII, CN VIII, and CN X. These pathways were tracked according to previous microdissection studies (Fernandez-Miranda et al., 2015; Wang et al., 2016). The course of the posterior column sensory pathway, running within the fascicles gracile and cuneatus toward the primary sensory cortex, was manually terminated at the level of the thalamus and labeled as ML. This segment in the brainstem corresponds to the second order neurons running from the nucleus gracile and cuneatus to the thalamus.

### Connectivity matrix, connectogram, and network measures

A weighted connectivity matrix was quantified using a cortical parcellation based on regions derived from the AAL atlas (Table S2). It is noteworthy that our tractography atlas can be readily applied to any cortical parcellation atlas, and currently there is no consensus on how network nodes should be defined. Here we used only one of the most popular parcellation from the AAL atlas to illustrate the network characteristics.

The average of along-track SDF values was used as the connectivity value. The connectograms of each fiber bundle and whole brain tracks were generated using CIRCOS (http://mkweb.bcgsc.ca/tableviewer/visualize/). The network measures such as network characteristic path length, global efficiency, local efficiency, clustering coefficient were calculated using the definition formulated in Brain Connectivity Toolbox (https://sites.google.com/site/bctnet/). The influence of the projection, association, and commissural pathways was calculated by calculating the change of network measures (quantified by percentage of the original) after removing the tracks.

The University of Pittsburgh Institutional Review Board reviewed and approved the study by the expedited review procedure authorized under 45 CFR 46.110 and 21 CFR 56.11 (IRB#: PRO16080387).

### Data and Code Availability

The processing pipeline (DSI Studio), SDF data of all 842 subjects, and HCP-842 template are available at http://dsi-studio.labsolver.org. The SDF template can be reproduced using the HCP data and documentation on the website. The atlas data, including the track trajectories and connectograms, are available at http://brain.labsolver.org.

## Results

### A high spatial and angular resolution diffusion template of the human brain

Diffusion MRI data from 842 participants were reconstructed in a standard space to calculate the SDF(Yeh and Tseng, 2011; Yeh et al., 2010) within each voxel (Fig. 1a). The goodness of fit between the normalized image and the template was reported as an R^2^ (Fig. S1). These values ranged from 0.73 to 0.86, and the quantiles were 0.81 (25%), 0.82 (50%), and 0.83 (75%), suggesting that the distribution of R^2^ values were mostly centered around 0.82, and more than 75% subjects had R^2^ values greater than 0.80. An SDF is an empirical distribution of the density of diffusing water orientations, calculated for each voxel to reveal the underlying fiber architectures (Fig. 2a). The SDFs of all subjects were averaged to build the HCP-842 SDF template, which represents an average diffusion pattern within a normal population (Fig. 1b and Fig. 2a). Figure 2b shows the peak orientations of fibers in each voxel, resolved from the group-averaged SDFs, near the corpus callosum crossing at central semiovale (red: left-right, green: anterior-posterior, blue: inferior-superior). The SDF peaks reflect the local orientation of underlying fiber bundles, whereas the magnitudes measured at the peaks provide estimates of the density of each bundle, which is used by the tractography algorithm determine the whether or not to terminate the tracking process. These two features offer the necessary information for a fiber-tracking algorithm to delineate long-distance white matter trajectories.

**Fig. 1.**
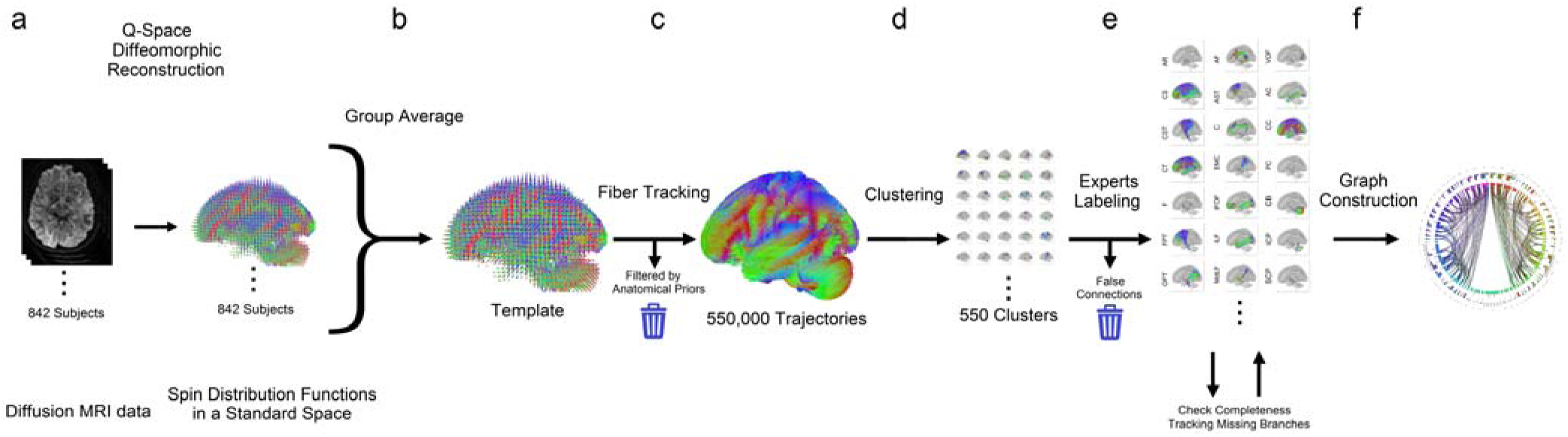
Flow chart of the processing steps used to construct a population-averaged structural connectome of the human brain. (a) A total of 842 subjects’ diffusion MRI data were reconstructed in a common standard space to calculate the spin distribution function at each imaging voxel. (b) The spin distribution functions were averaged to build a template of the diffusion characteristics of the normal population. (c) The template was used to guide a fiber tracking algorithm and generate a total of 550,000 trajectories. (d) Automatic track clustering was applied to cluster trajectories into fiber bundles. (e) A team of experienced neuroanatomists manually labeled each cluster and identified false pathways according to the neuroanatomy evidence. The clusters with the same labeled were grouped together as an atlas of structural connectome. An additional quality check was conducted to ensure complete coverage. (f) The atlas was then used to build the connectogram showing the connections between brain regions.

**Fig. 2.**
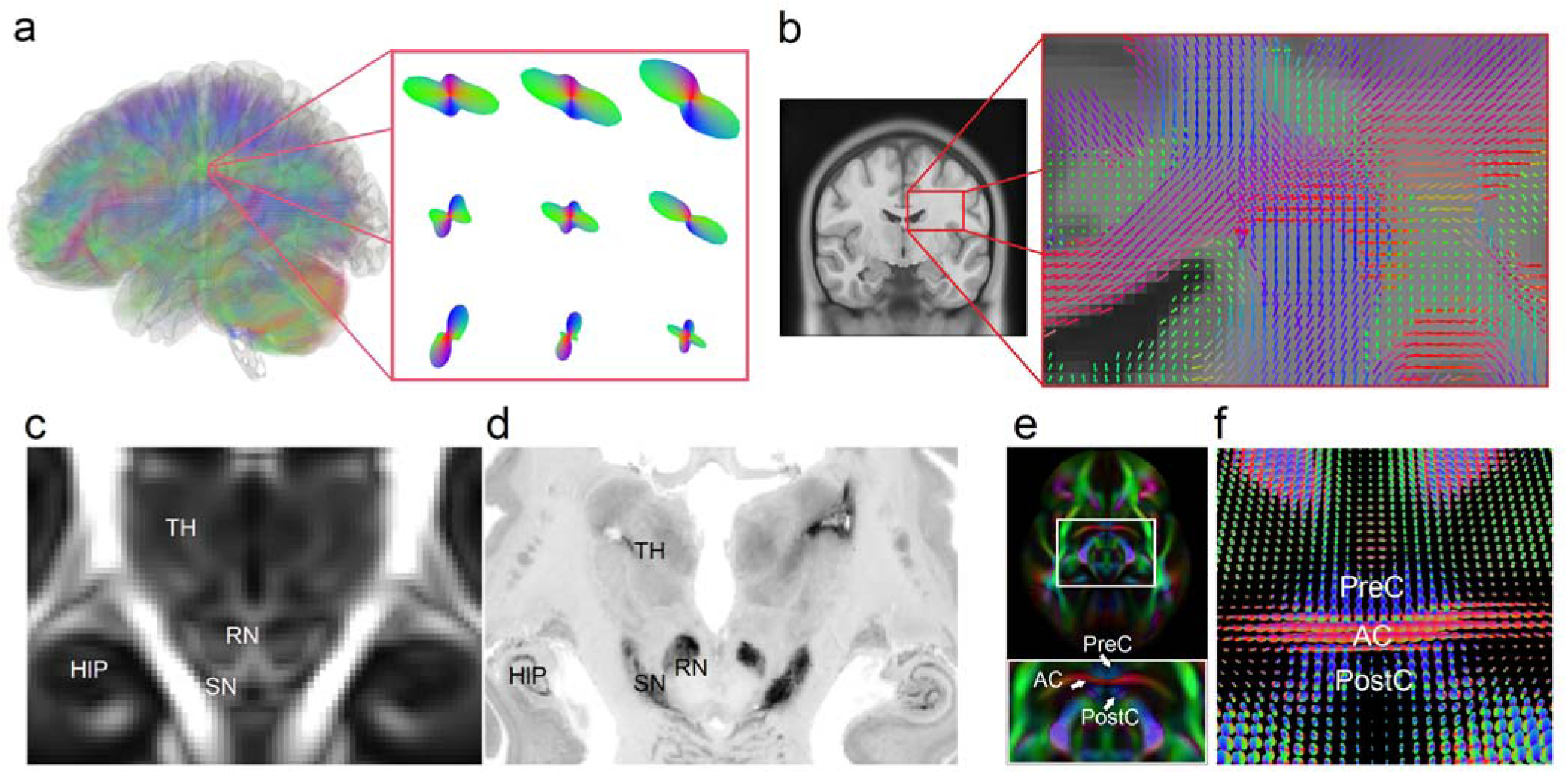
(a) Diffusion MRI allows for quantifying, for each imaging voxel, the orientation distribution of the water diffusion (termed spin distribution function, SDF) to reveal the underlying structural characteristics of axonal fiber bundles in a color-coded surface (red-blue-green indicates the orientation at the x-y-z axis, respectively). The protruding points of the SDFs indicate the orientation of fiber bundles. (b) The color sticks represent the peak orientations on SDFs. The coronal view shows that SDF can resolve crossing fibers at central semiovale, a white matter region where the corpus callosum crosses vertical passing fibers. The SDFs averaged from a total 842 subjects provide orientations of the local axonal connections. The information can be used to drive a fiber tracking algorithm to delineate white matter connections. (c) The SDF template of the human brain averaged from 842 diffusion MRI scans (termed the HCP-842 template) shows structural characteristics of the human brain. The magnitude map of the HCP-842 template reveals structures such as hippocampus (HIP), thalamus (TH), red nucleus (RN), and substantia nigra (SN), which are consistent with the histology image from BigBrain slides (d). (e) The orientation map of the HCP-842 template allows for delineating the complicated structures, such as the clamping structure between the anterior commissures (AC) and the pre-commissural (PreC) and post-commissural (PostC) branches of the fornix. The structural characteristics are also illustrated by the SDFs of the HCP-842 template in (f).

Although the group-averaged SDFs appear smoother due to the averaging effect, they are still capable of resolving major crossing architectures. The number and percentage of voxels that contain more than one fiber orientations are listed in Table 1. These results were obtained by re-gridding the template at different resolutions to aggregate information about underlying fiber pathways in each voxel. The table shows that after re-gridding at 2-mm^3^ and 2.5-mm^3^ resolution, more than 80% of the white-matter voxels in the HCP-842 template had more than one distinct fiber orientation. These percentage values are consistent with previous estimates of 60~90% of voxels having multiple fiber orientations when sampled and reconstructed at a 2.4mm^3^ resolution (Jeurissen et al., 2013). It is noteworthy that the percentage of multi-fiber voxels dropped substantially at 1.5-mm^3^ and 1-mm^3^ resolutions. This can be explained by the fact that at a smaller voxel sizes, turning, and branching configurations can be spatially resolved into one single fiber orientation and the number of voxels containing multiple fiber orientations drops dramatically.

**Table 1.** **Number of voxels with more than on fiber orientations resolved in different resolutions**

| Resolution | Voxel with more than one<br>fiber orientations* | Total white matter voxels | Percentage (%)* |
| --- | --- | --- | --- |
| 1 mm | 76452 | 606662 | 12.60 |
| 1.5 mm | 114856 | 195705 | 58.69 |
| 2 mm | 74283 | 89306 | 83.18 |
| 2.5 mm | 43551 | 49050 | 88.789 |

\* may include turning, crossing, or branching fibers

Qualitatively, the HCP-842 appears to resolve underlying neuroanatomical architecture with high fidelity in spatial resolution. Comparing a coronal slice of the HCP-842 (1-mm resolution, Fig. 2c) with a similar section from the BigBrain histology image (the 200-micron resolution version, Fig. 2d), we see that HCP-842 clearly delineates subcortical structures such as the hippocampus (HIP), substantia nigra (SN), red nucleus (RN), and thalamus (TH). The high spatial resolution of the orientation map is even more apparent at the anterior commissure (AC) (Fig. 2e), a small left-right connecting pathway clamped by the pre-commissural (PreC) and post-commissural (PostC) branches of fornix that run in the vertical direction (color-coded by blue). The clamping structure formed between AC and fornix is a benchmark for examining the spatial resolution of the template. Figure 2e resolves AC from the PreC and PostC branches, whereas Figure 2f shows the averaged SDFs at the same region depicting the structural characteristics of AC with the PreC and PostC branches of the fornix. The ability to resolve branches of fornix from AC reveals the intricate sensitivity of the HCP-842 to map detailed brain connections.

### Supervised labeling and segmentation of major pathways

To isolate major and minor white matter fascicles, we applied whole-brain fiber tracking to the HCP-842 group-average template, producing a total of 550,000 fiber trajectories in the standard space to achieve an average density of 1 track per voxel (Fig. 1c). A white matter mask was used to remove tracks that have premature terminations in the core white matter. The remaining whole-brain tracks were then automatically clustered by a single-linkage clustering algorithm, generating unique clusters of fiber bundles (Fig. 1d). The trajectories that were proximally close to one another were grouped. Each cluster could subsequently contain a different number of trajectories based on the anatomical proximity of the tracks. Figure 3 shows the largest 40 out of the 550 clusters as an example, where the size of a cluster is determined by the number of its containing tracks. Shorter pathways, such as the uncinate fasciculus, will receive less seeding counts in the tracking process and thus be estimated to have a smaller size. Many track bundles were also represented by more than one cluster component. For example, cluster #1 and #38 are both labeled as the corpus callosum. A team of clinical neuroanatomists then examined and labeled the clusters according to neuroanatomical nomenclature. Table S1 lists all labels used in naming the clusters and the relevant neuroanatomy literature used for examination. Label “X” indicates a false track, which may arise due to false continuations (Fig. 4a) or premature termination (Fig. 4b). Only the 550 largest clusters were used because the false rate (either false continuation or premature termination) increased substantially in clusters with a smaller size (Fig. 4c). The labeled clusters were subsequently merged according to their neuroanatomical label. Missing components of the large fiber bundles were tracked separately and merged to ensure completeness as per the literature (Fig. 1e).

**Fig. 3.**
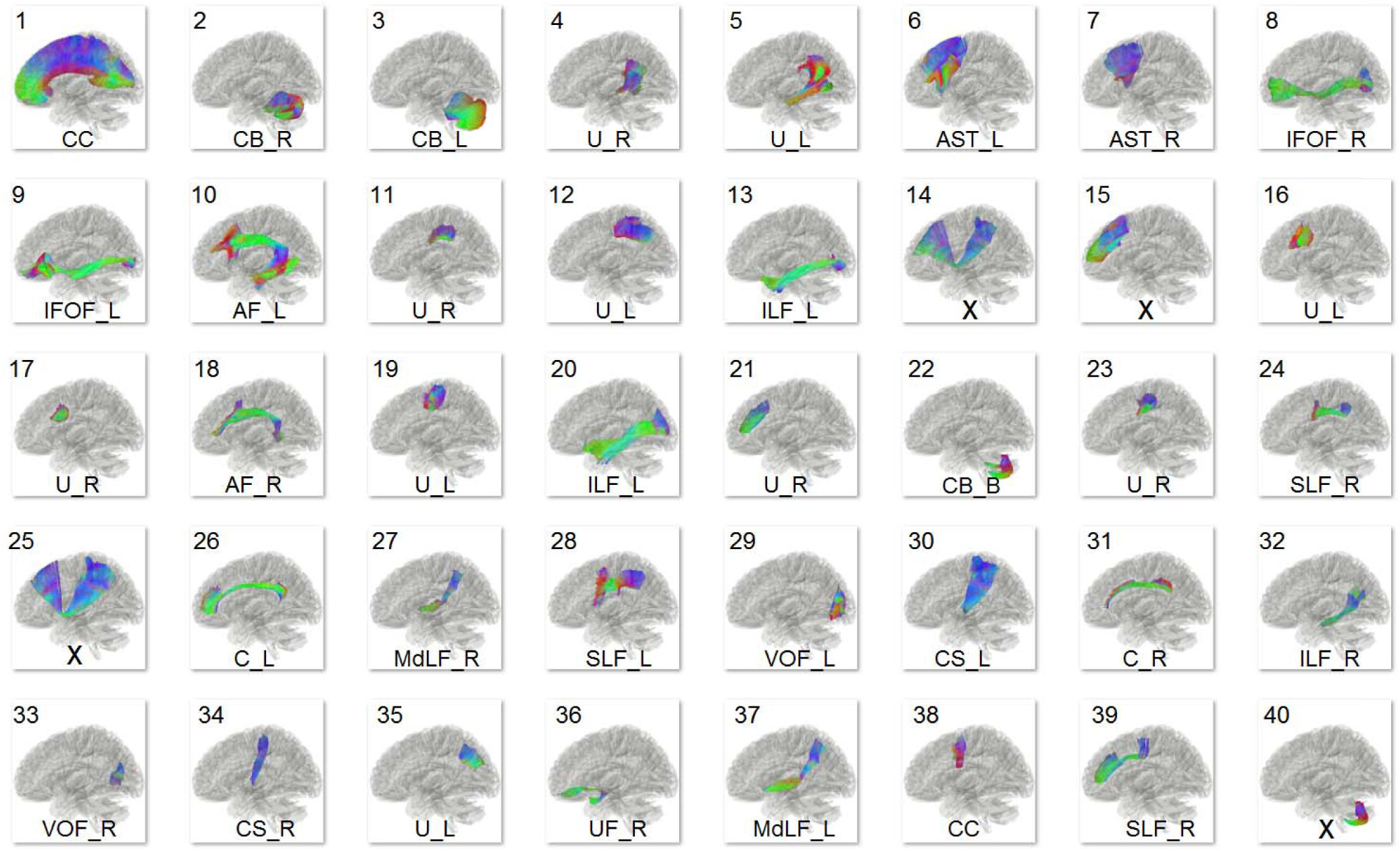
The 40 largest clusters (selected from a total of 550 clusters) generated from automatic track clustering and their labels assigned by neuroanatomists. False connections are assigned by “X”, whereas the others assigned by their corresponding neuroanatomy abbreviations.

**Fig. 4.**
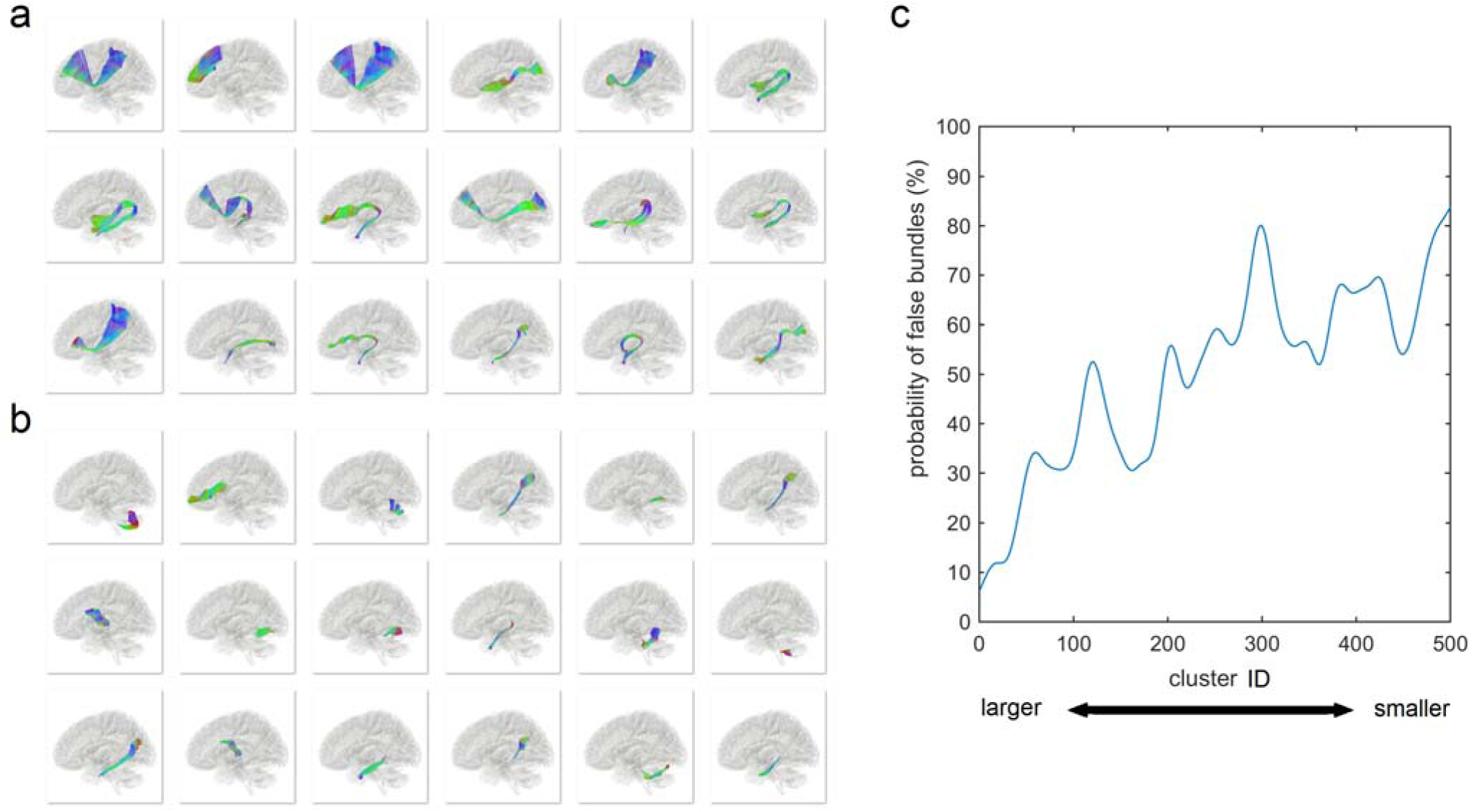
False connections due to (a) false continuation and (b) premature termination identified by the neuroanatomists. A false continuation is a common cause of false trajectories and often found in regions with two fiber population cross on top of each other. Premature termination is often due to a failure in resolving crossing or branching pattern in the white matter. (c) The probability of a cluster labeled as “false” increases substantially with decreased cluster size. This suggests we can discard smaller clusters as there are mostly false connections.

The high-angular-resolution quality of the atlas can be appreciated in the corticospinal and corticobulbar tracts generated from our pipeline (Fig. 5a). These show a fanning projection pathway from the precentral (motor) cortex along the cortical surface that is consistent with the anatomical evidence (right, modified from Gray’s Anatomy). In addition, the coronal view of the corpus callosum (Fig. 5b) also shows a widespread fanning pattern, not otherwise trackable using lower angular resolution methods. The midline portion of the corpus callosum tracks (Fig. 5c) shows matching volume with the ICBM-152 T1-weighted images, suggesting that the atlas can also provide volumetric measurements. Thus, the atlas appears to capture more complete portions of major pathways that are typically lost using traditional approaches.

**Fig. 5.**
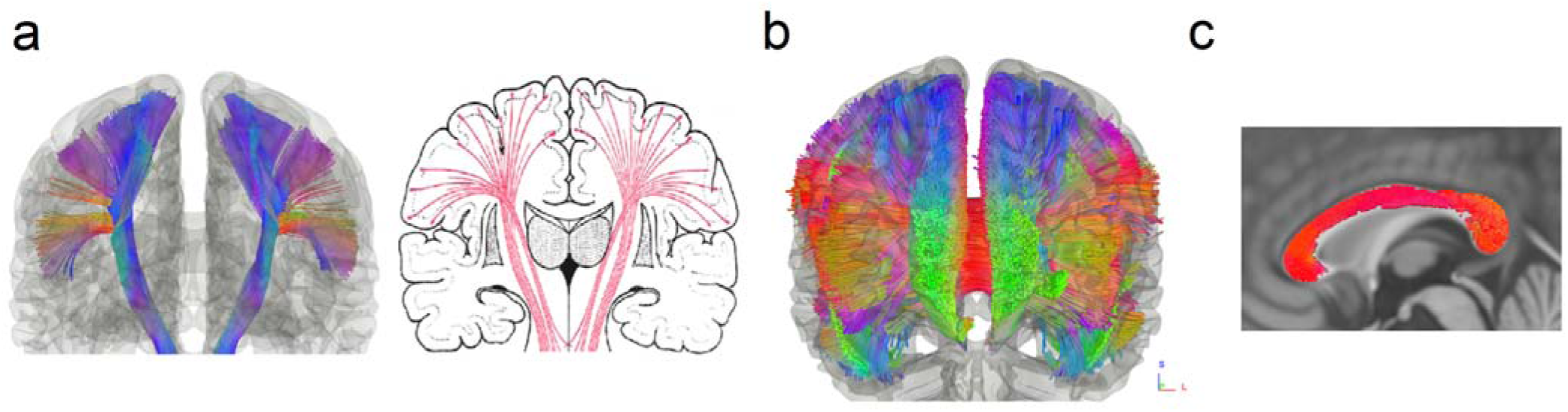
The angular resolution of the structure connectome atlas illustrated. (a) The corticospinal and corticobulbar tracks in the structure connectome atlas present a fanning pattern consistent with the known neuroanatomy presentation. (b) The coronal view of the corpus callosum mapped by the atlas shows a wide spreading fanning pattern, which cannot be achieved using a low angular resolution approach. (c) The mid portion of the corpus callosum matches well with the ICBM-152 T1-weighted images, suggesting that the track bundles in the atlas have volumes matching the standard template.

### A population-averaged atlas of macroscopic structural connectome

The full atlas of the structural connectome is shown in Fig. 6 (abbreviation listed in Table S1) and includes the most comprehensive map of white matter pathways yet reported. This includes the projection pathways that connect cortical areas with subcortical nuclei and brainstem. Acoustic radiation has not been previously reported in tractography due to the complicated crossing pattern of the pathway. The association pathways connect disparate cortical areas, including a set of U-fibers (U). The commissural pathways connect the two hemispheres and include the corpus callosum, anterior commissure, and posterior commissure. Posterior commissure has not been previously reported in tractography. The cerebellar pathways include the cerebellar tracts (CB) and peduncles (SCP, MCP, ICP), and they provide the major input, output, and internal connectivity of the cerebellum. We were even able to resolve several brainstem pathways, such as central tegmental tract (CTT), dorsal longitudinal fasciculus (DLF), lateral lemniscus (LL). Finally, we discovered a limit of the current spatial resolution, where a set of cranial nerves including CN III, CN VII, and CN VIII were successfully identified, but CN I, IV, VI, and IX could not be identified due to insufficient spatial resolution. The detailed connective routes of the structural connectome atlas are presented in Supporting Information, including projection pathways (Fig. S2), association pathways (Fig. S3), commissural pathways (Fig. S4), cerebellar pathways (Fig. S5), brainstem pathways (Fig. S6), and cranial nerves (Fig. S7). It is worth noting that several cranial nerves cannot be found in the HCP-842 template due to the limitation of its spatial resolution. The full atlas, including the track trajectories and connectograms, is publicly available at http://brain.labsolver.org.

**Fig. 6.**
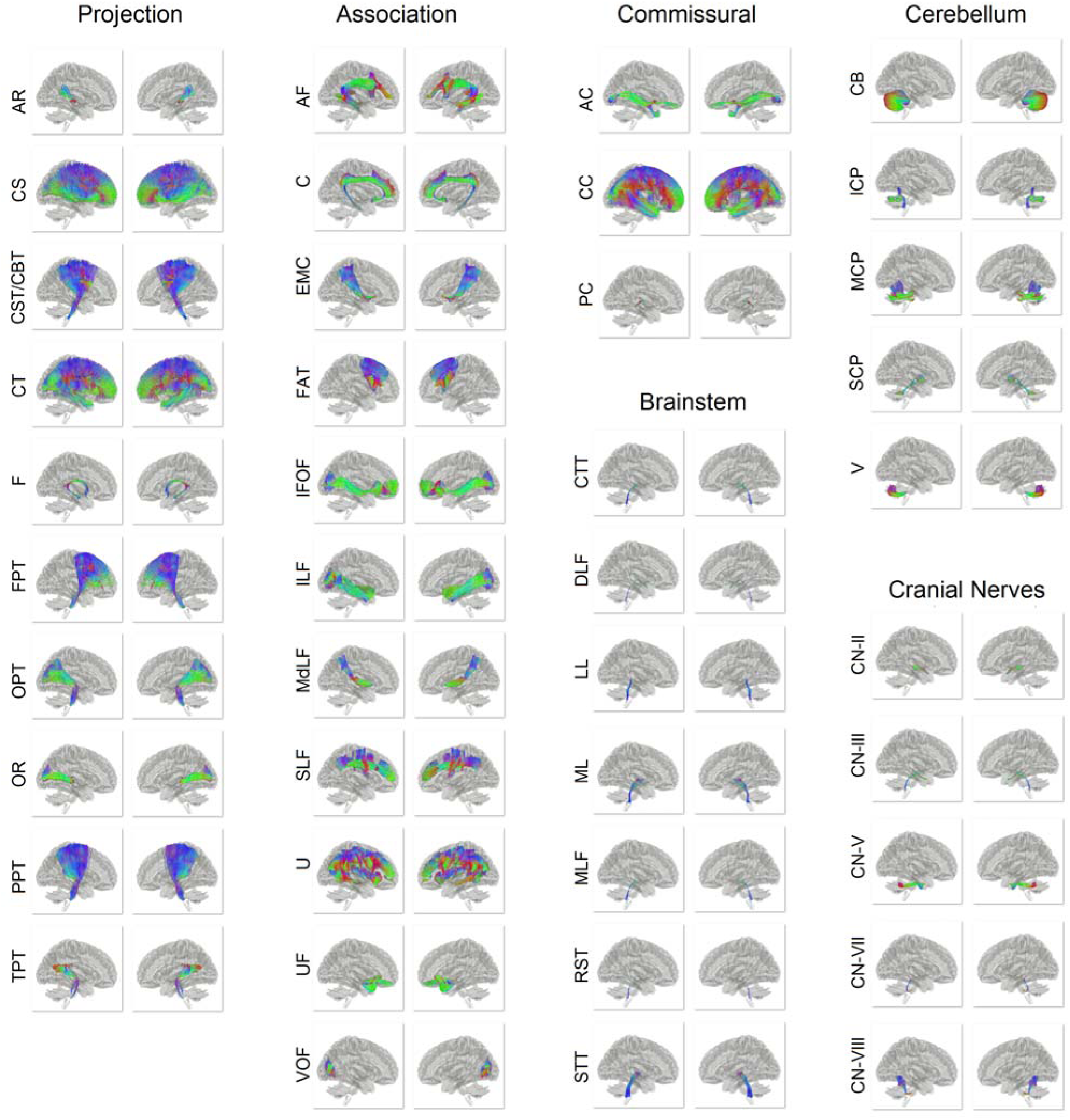
Overview of the population-averaged structural connectome atlas categorized into the projection, association, and commissural pathways in addition to cerebellum pathways, brainstem pathways, and cranial nerves. Each pathway contains thousands of trajectories showing the representative connections of the 842 subjects between brain regions in a standard space. The trajectories are color-coded by the local orientation (red: left-right, green: anterior-posterior, blue: inferior-superior). This connectome atlas provides normative connection routes between brain regions that can facilitate network analysis, simulation and modeling.

### Neuroanatomical constraints on connective topology

The atlas of the structural connectome from the HCP-842 addresses a critical need in connectivity estimates that suffer from a high false positive error rate (Maier-Hein et al., 2016; Thomas et al., 2014): the atlas enables estimation of normative region-to-region connectivity that is anatomically constrained. Figure 7 shows region-to-region connectivity matrix weighted by the SDF magnitude along the fiber pathways, segmented into the projection, association, and commissural pathways. The abbreviations for brain region are listed in Table S2. Higher intensity (white) indicates greater SDF magnitude along the pathway. This anatomically-constrained view of structural connectivity between gray matter targets highlights how specific classes of white matter pathways define unique connective topologies. For example, commissural pathways have a generally symmetrical topology of connections between the hemispheres, with greater homotopic connectivity than heterotopic connectivity, whereas the association pathways are more uniform in their intra-cortical connections.

**Fig. 7.**
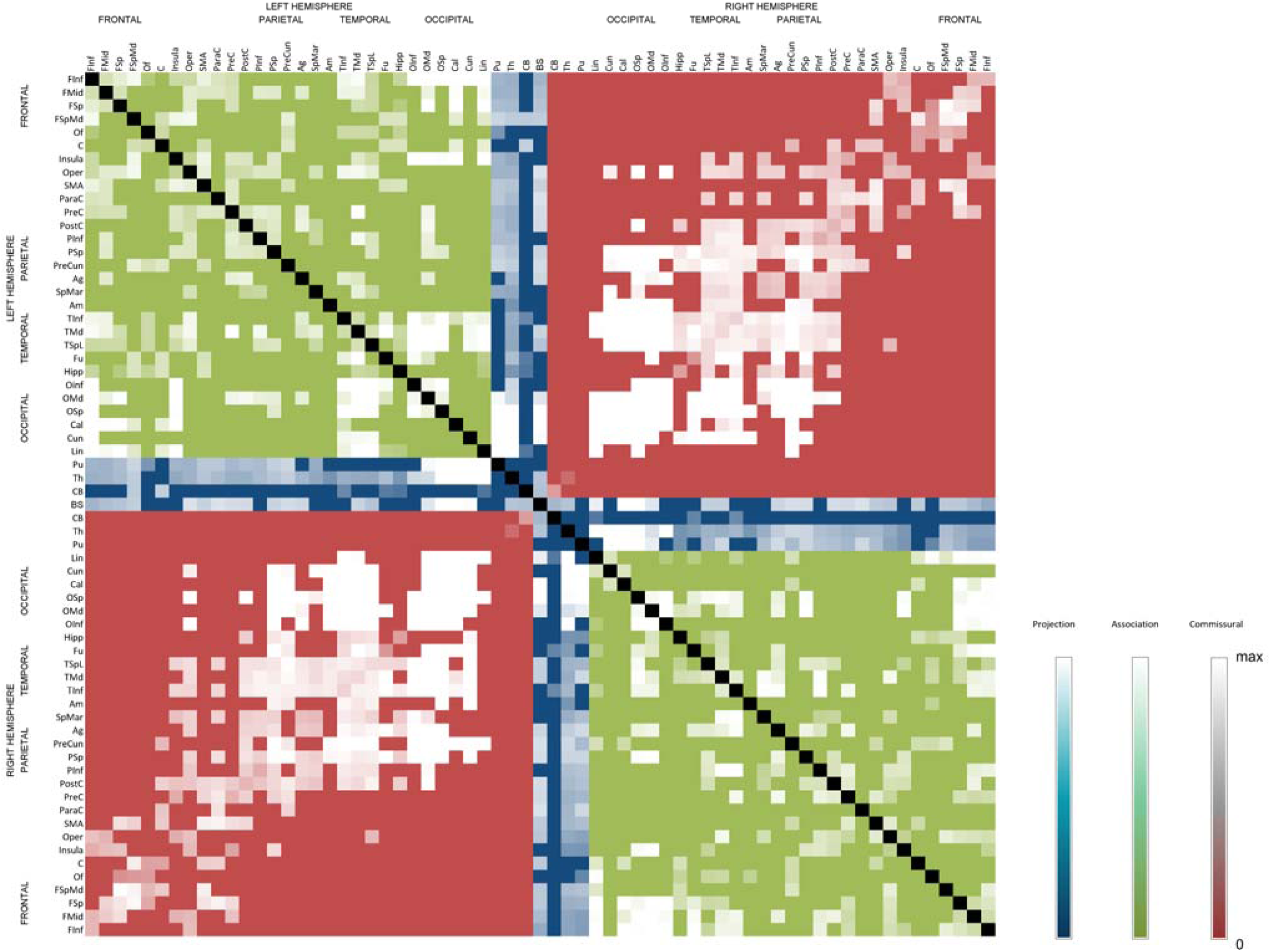
The connectivity matrix constructed from the human connectome atlas. The color division shows the division of three major track systems—projection (blue), association (green), and commissural (red)—in the human brain. The intensity shows the between region connectivity quantified the magnitude of the along-track diffusion properties quantified by spin distribution functions.

Finally, the connectograms of the structural connectome are illustrated in a multi-level approach (Fig. 8). The connectogram of the whole brain pathways illustrates the first level of the gross network topology (Fig. 8a, the high-resolution version shown in Fig. S8). The overall figure shows a dense network topology, and its network characteristics cannot be readily visualized due to the high complexity of the brain network at this level. The connectograms of the projection, association, and commissural pathways in Fig. 8b, 8c, and 8d depict the second level of the network topology (high-resolution details in Fig. S9), and within this level, the connectograms start to reveal important network features. The projection pathway in Fig. 8b indicates hub structures at thalamus, putamen, and brainstem, illustrating the role of these regions in integrative sensorimotor function between the cerebral cortex and corresponding peripheral systems. The association pathway, as shown in Fig. 8c, forms clusters within each hemisphere and contributes a substantial amount of clustering coefficient and local efficiency (Table 2), elucidating its small-worldness that involves multiple relevant gray matter regions. The commissural pathways, as shown in Fig. 8d, serve as a bridge connecting both hemispheres and provide global efficiency (Table 2) to integrate information across cerebral hemispheres. In Fig. 8e, the connectograms of each fiber bundle are further divided to show the third level of the network topology in much more detail, and the illustration reveals a consistent hub formation for different fiber bundles, albeit with an alternative connectivity pattern to the cerebral cortex. Fig. 8f also shows clustering topology within different cortical areas, whereas Fig. 8g shows bridge-like symmetric structures of inter-hemisphere connections. Together, these unique topologies based on the class of fiber pathway highlights the rich taxonomy of structural connectome in the human brain that reflects unique information processing constraints.

**Fig. 8.**
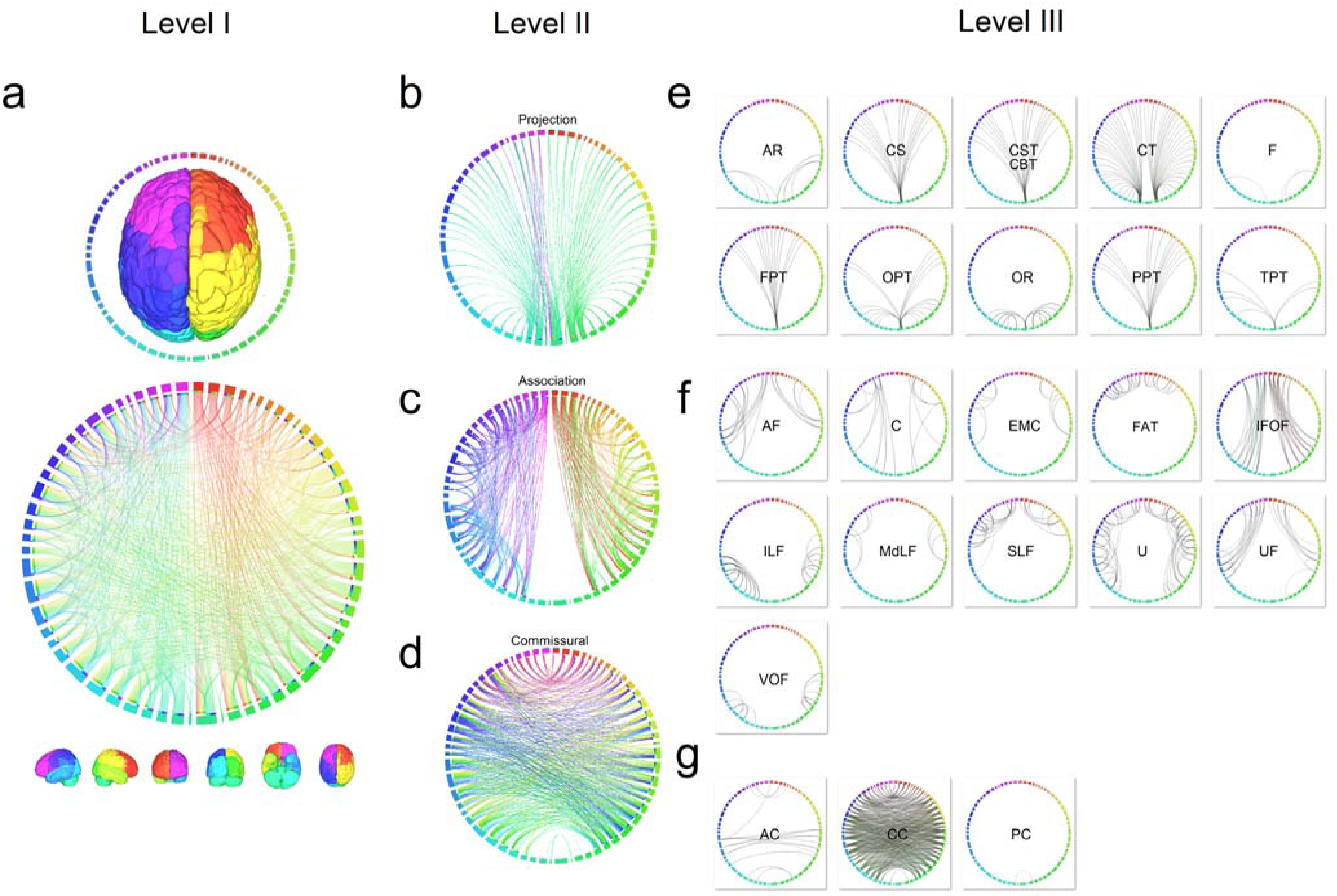
The multi-level connectograms of the human structural connectome. (a) The first level of the overall structural connectome shows a dense connections pattern in the average structure connectome. (b) The second level of the connectogram shows the network characteristics in each pathway system. The projection pathway forms a hub structure at thalamus, putamen, and brainstem. The association pathway is constituted of numerous clusters in the brain networks. The commissural pathway has long-range connections between hemispheres that provide global efficiency. (c) The third level of the connectogram reveals the network pattern of each fiber pathways under the projection, association, and commissural system. The connection patterns inherit the characteristics of their belonging pathway system shown in the second level connectogram.

**Table 2:** **Change of network measures with/without projection, association, and commissural pathways**

| Pathway<br>System | Clustering<br>Coefficient (%) | Network |  |  |  |
| --- | --- | --- | --- | --- | --- |
|  |  | Characteristic<br>Path Length<br>(%) | Global<br>Efficiency (%) | Local<br>Efficiency (%) | Small<br>Worldness (%) |
| Projection | -0.30 | -5.38 | 3.35 | 4.60 | 5.08 |
| Association | 62.27 | -7.95 | 6.88 | 57.90 | 66.91 |
| Commissural | -18.65 | -10.57 | 8.11 | -7.47 | -8.24 |

A positive value indicates an increase of network measures *with* the pathways added.

## Discussion

Here we present the first complete population-level atlas of the human structural connectome and its network topology, delineating fiber pathways within the cerebrum, cerebellum, brainstem, and a subset of cranial nerves. The fiber trajectories were generated from a group-averaged template of 842 subjects using a fiber tracking algorithm that has been shown to minimize tracking errors relative to other methods (Maier-Hein, 2016). Using an automated clustering approach, tracks were grouped into small bundles and subsequently labeled by a team of clinical neuroanatomists and vetted according to their neuroanatomic nomenclature. This combination of optimizing strategies allowed us to construct a high-quality, group-averaged structural connectome atlas of the human brain, and this HCP-842 atlas and its associated data set will be made publicly available (http://brain.labsolver.org) to promote future connectomic studies and assist neuroscientists to gain insight into the structural topology of the human brain.

We should note that several human white matter atlases have been previously released. These include voxel segmentations on individual subjects that label the core of major pathways (Mori et al., 2009; Mori et al., 2008; Peng et al., 2009; Zhang et al., 2011) or tractography atlases based on tracking individual subjects data (Catani et al., 2002; Guevara et al., 2012; Thiebaut de Schotten et al., 2011; Zhang et al., 2008).Our atlas expands on these currently available resources by providing a comprehensive characterization of normative major and minor white matter fascicles constructed from a large sample of 842 individuals who were imaged using high angular and high spatial resolution diffusion acquisitions, allowing for the resolution of multiple fiber populations within a white matter region to delineate the intertwining architecture of human white matter. This novel population-level description of the structural connectome characterizes both the normative 3D trajectories of white matter fascicles and delineates *how* gray matter regions in the cerebrum, cerebellum, and brainstem are physically connected by nearly all macroscopic white matter pathways. For the first time, this atlas offers structural detail and network topology of both large and small pathways, such as the clamping structure between the fornix and the anterior commissure that cannot be discerned from individual studies due to lower resolution and signal-to-noise ratio of conventional diffusion MRI.

While overcoming many challenges, our current approach still has its limitations. First, our atlas does not address the variability of the fiber pathways across subjects. While it is entirely feasible to repeat the fiber tracking procedures for each of the HCP subjects, the labeling of 550 clusters of all 842 subjects may require a substantial amount of expert efforts. This labor-intensive approach would require several years worth of human labor to complete. Thus an automated approach to replace expert labeling would better assist this future endeavor, but developing such an automated classifier is well beyond the scope of the current study. In addition to individual variability, there could be errors in manual labeling of the clusters, and thus there should be better ways to address the inter-observer differences, as we only resolved differences by a group discussion with the goal of reaching a consensus on every track. This would save some time, but not enough to make this feasible and extensible enough to use in applied studies. Of course, there are also controversies in neuroanatomical structures (Meola et al., 2015) that can be further complicated by individual differences. Thus, we have made all of the clusters data, their labels, and the entire atlas publicly available, thereby allowing for future modifications to improve the atlas as well as the development of better tools for automated segmentation.

Moreover, the fiber tracking algorithm used in this study could still have false positive and false negative results. While expert assistance may address part of this issue, it cannot handle the false negative problem, and there could be missing tracks in our atlas. For example, several cranial nerves that are smaller than 1-mm in width were not detected by our method. These can only be tracked using images acquired at a much higher resolution. In addition, the expert examination may have its own errors, especially for identifying minor pathways and branches. It is also possible that the branching patterns of the white matter pathways differ person from person. Finally, the atlas reveals only three levels of the network topology, as more recent studies have focused on detailed subcomponents of the fiber bundles (e.g. SLF I, II, and III)(Fernandez-Miranda et al., 2015; Wang et al., 2016). Although the spatial resolution of the atlas can be improved, it provides a macroscopic framework for future connectomic studies to explore microscopic connections under its categorical system.

Despite these limitations, a vetted atlas of the population-level structural connectome has many benefits for clinical, scientific, and educational applications. The atlas can be used to derive a representative pattern of network measures to assist graph theoretical analysis of clusters and hubs in the brain connectome. It can be used to confirm or explore potential cortical connections from functional measures (e.g., functional connectivity), augmenting current functional-structural correlative inferences or supplementing prior anatomical connectivity expectations in studies that do not have access to individual dMRI data. This, for example, may enable future investigations into the correlation of white-matter lesions with known gross-white matter structures. Another advantage of the current atlas is that it includes a normative template of diffusion distribution across the brain. This may allow for future efforts comparing normal diffusion patterns with those from the neurological or psychiatric pathologies. Finally, in science education, the atlas is a novel resource superseding conventional 2D slice-based histological atlases. The trajectory information provides panoramic views on the relative location of each white matter bundle, allowing for an in-depth understanding of the white matter structure.

## Acknowledgements

Data were provided in part by the Human Connectome Project, WU-Minn Consortium (D. Van Essen and K. Ugurbil, 1U54MH091657 NIH). This research was sponsored by the Army Research Laboratory and was accomplished under Cooperative Agreement Number W911NF-10-2-0022 and NSF BIGDATA grant #1247658. The views and conclusions contained in this document are those of the authors and should not be interpreted as representing the official policies, either expressed or implied, of the Army Research Laboratory or the U.S. Government. The U.S. Government is authorized to reproduce and distribute reprints for Government purposes notwithstanding any copyright notation herein.

## SUPPLEMENTARY MATERIALS

**Fig. S1.**
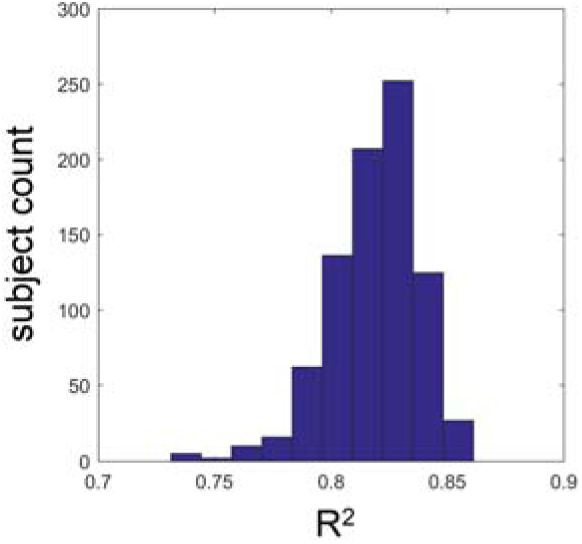
The histogram showing the empirical distribution of the goodness-of-fit values of the spatial normalization. The goodness-of-fit was quantified by the R^2^ value between each subject’s normalized anisotropy map and the template. The distribution shows a skewed function with a tail of lower goodness-of-fit values, which allowed us to examine datasets with potential quality issues.

**Fig. S2.**
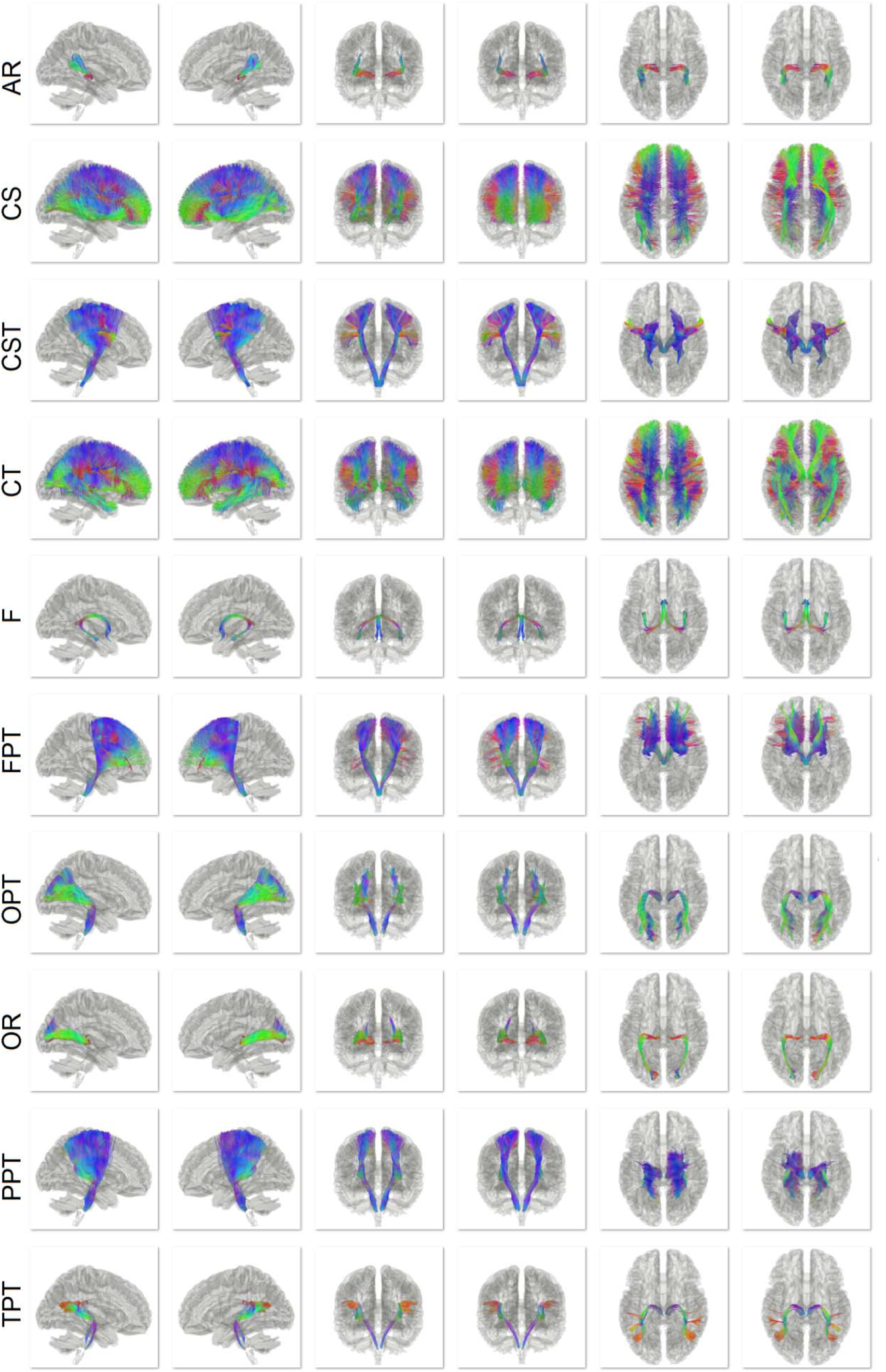
The fiber bundles in the projection pathways, including acoustic radiation (AR), corticostriatal pathway (CS), corticospinal tract (CST), corticothalamic pathway (CT), fornix (F), frontopontine tract(FPT), occipitopontine tract (OPT), optic radiation (OR), parietopontine tract (PPT), and temporopontine tract (TPT).

**Fig. S3.**
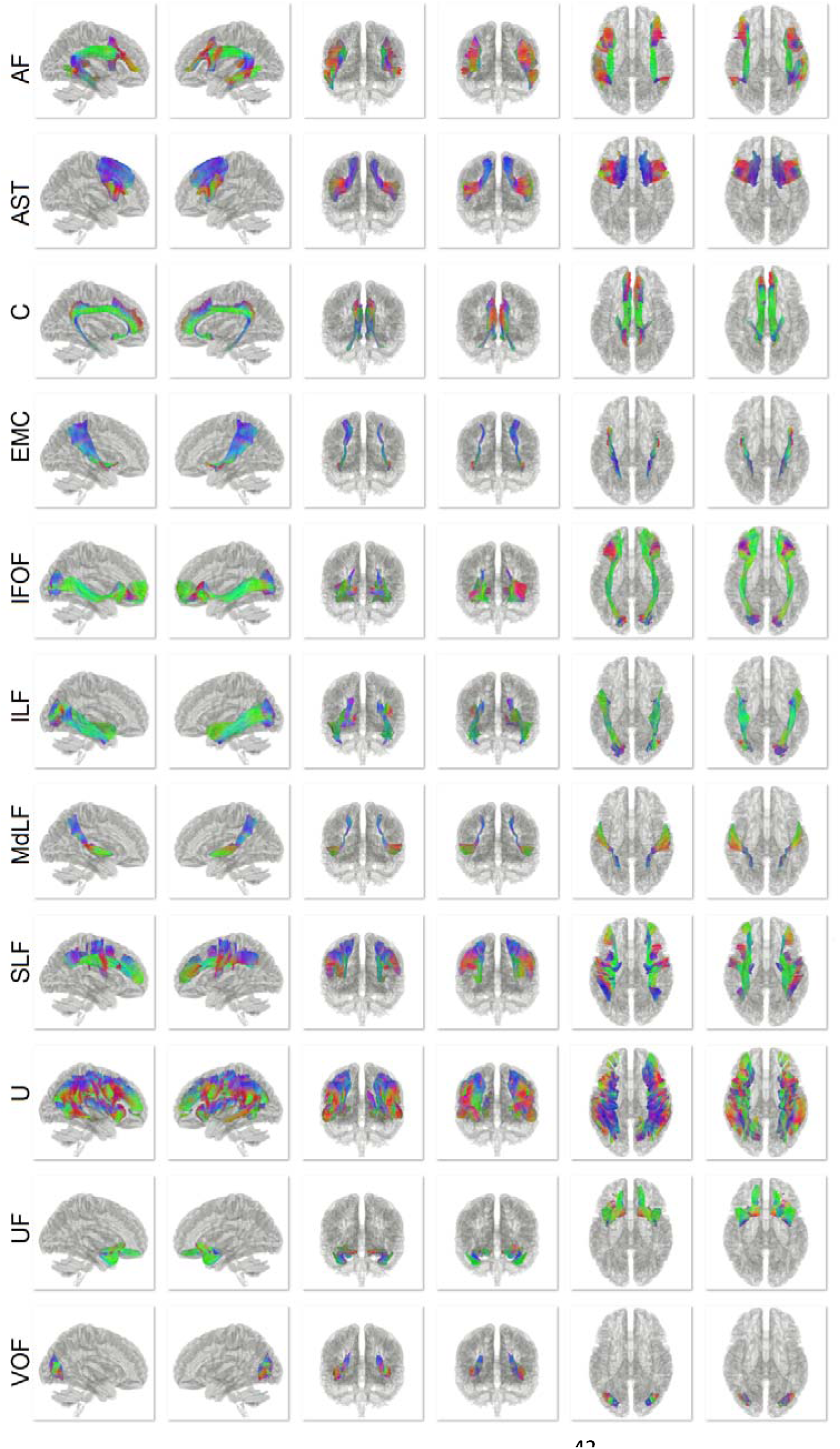
The fiber bundles in the association pathways, including arcuate fasciculus (AF), frontal aslant tract (AST), cingulum (C), extreme Capsule (EMC), inferior fronto-occipital fasciculus (IFOF), inferior longitudinal fasciculus (ILF), middle longitudinal fasciculus (MdLF), superior longitudinal fasciculus (SLF), U-fibers (U), uncinate fasciculus (UF), and vertical occipital fasciculus (VOF).

**Fig. S4.**
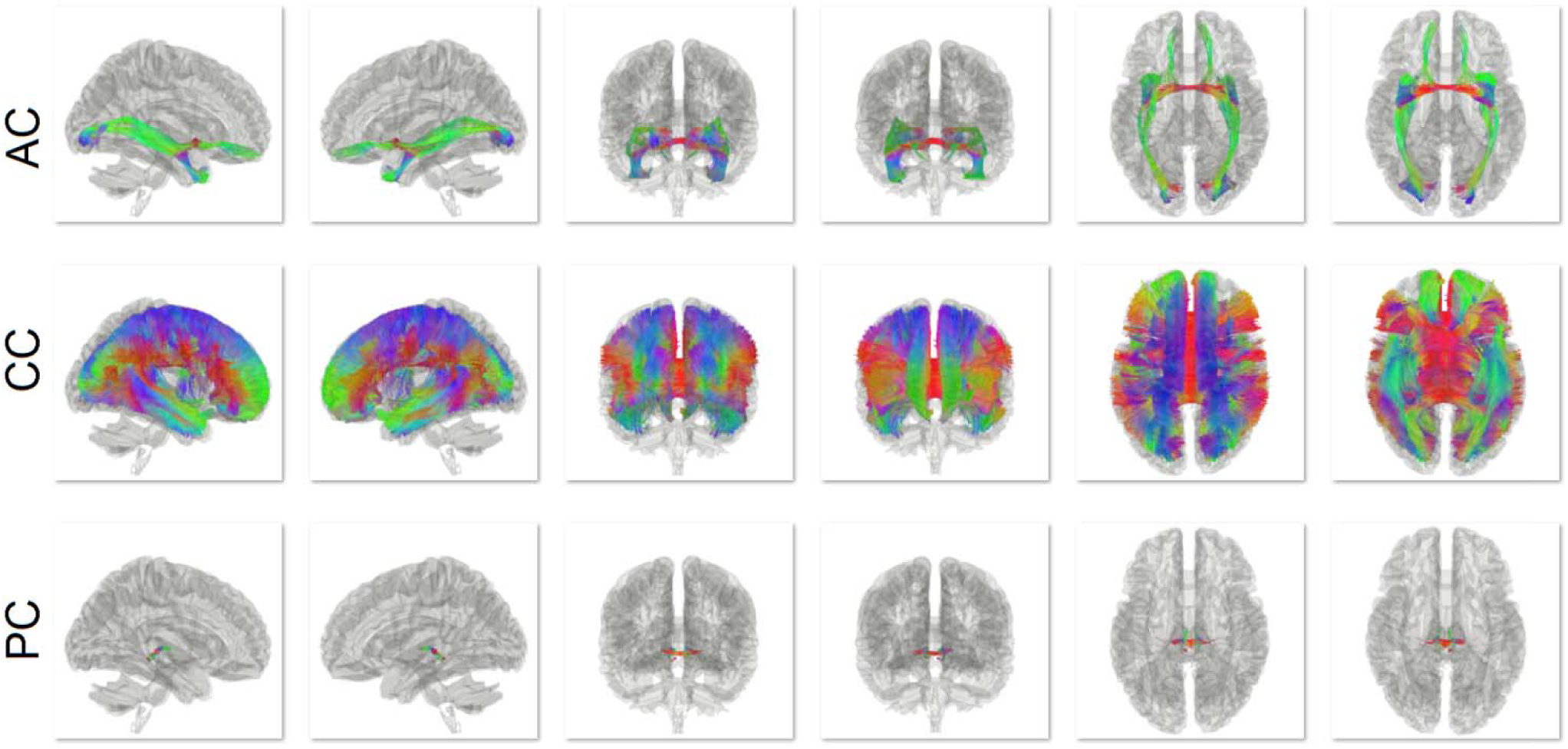
The fiber bundles in the commissural pathways, including the anterior commissure (AC), corpus callosum (CC), and posterior commissure (PC).

**Fig. S5.**
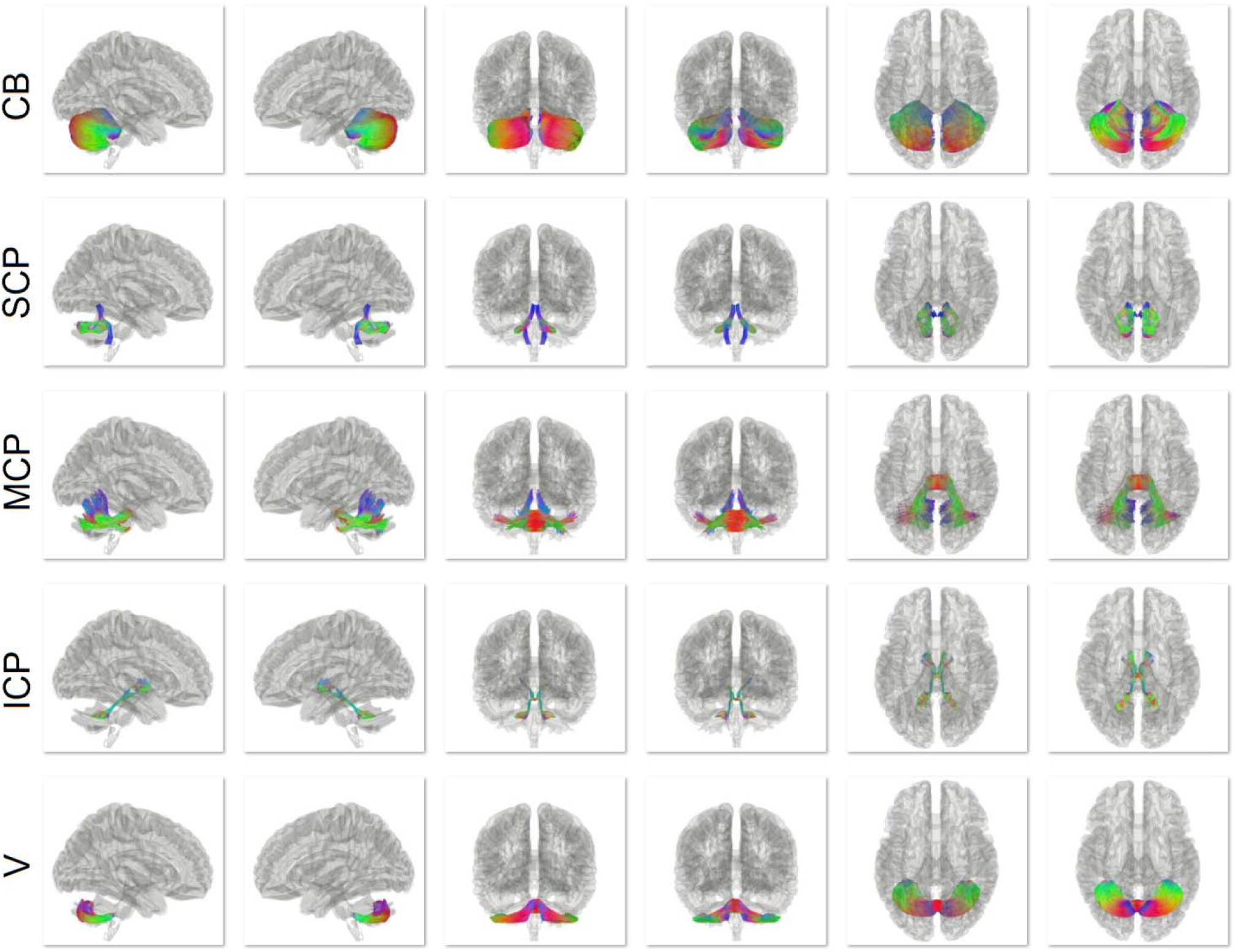
The fiber bundles in the cerebellar pathways, including the cerebellum (CB), superior cerebellar Peduncle (SCP), middle cerebellar peduncle (MCP), inferior cerebellar peduncle (ICP), and vermis (V)

**Fig. S6.**
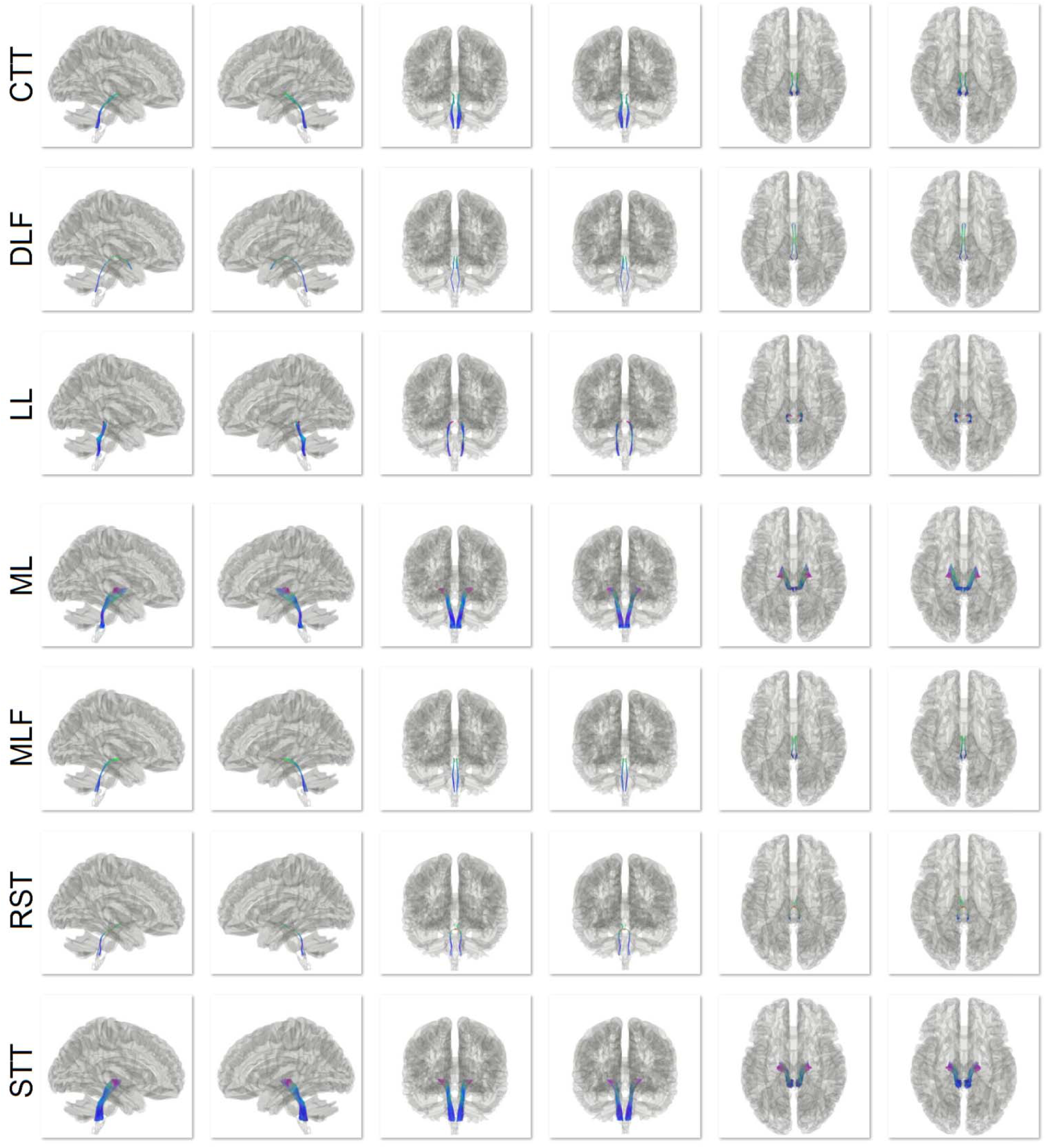
The fiber bundles in the brainstem, including central tegmental tract (CTT), dorsal longitudinal fasciculus (DLF), lateral lemniscus (LL), medial lemniscus (ML), medial longitudinal fasciculus (MLF), rubrospinal tract (RST), and spinothalamic tract (STT).

**Fig. S7.**
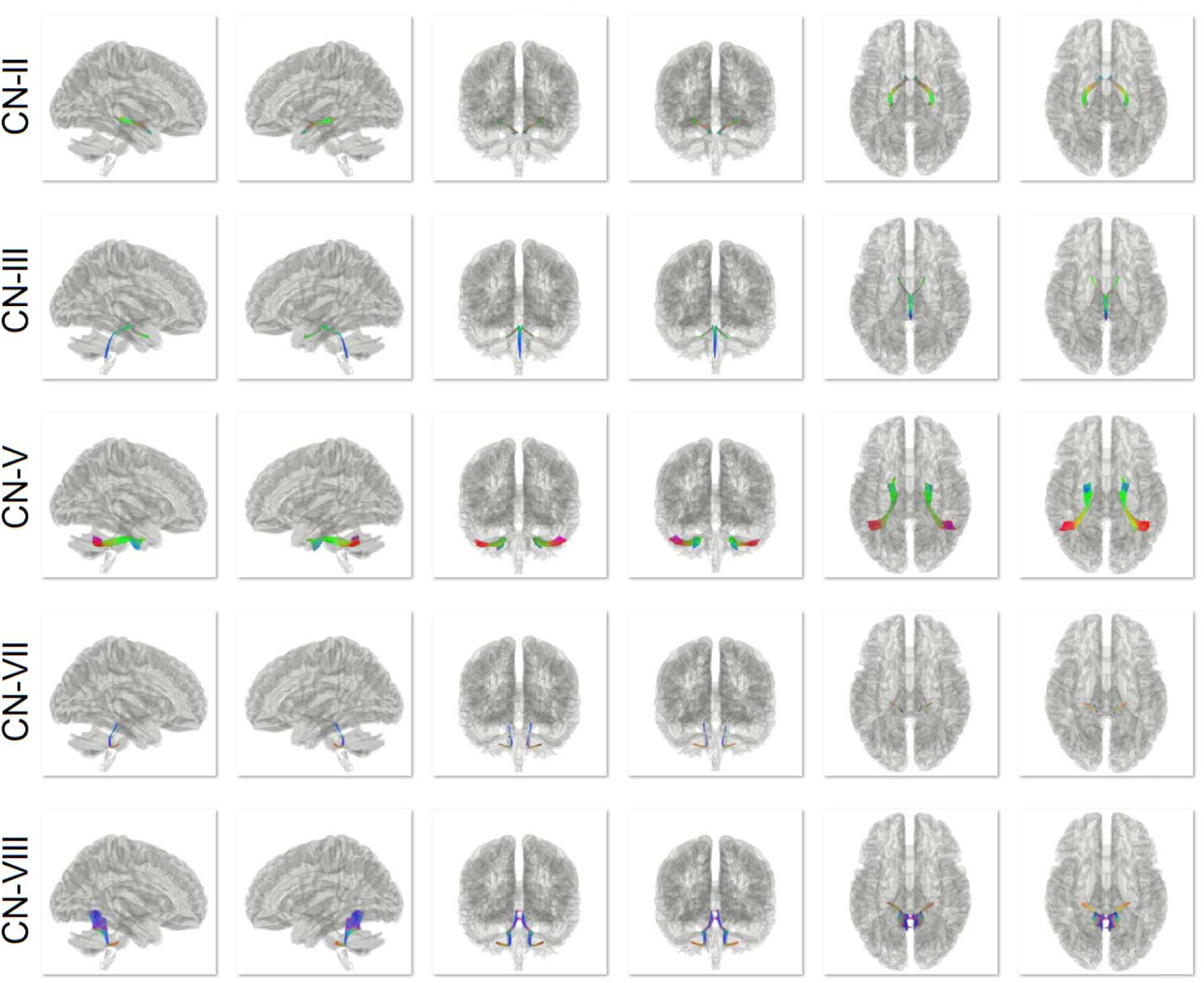
The cranial nerves included in the atlas, including the visual nerve (CN II), oculomotor (CN III), trigeminal nerve (CN V), facial nerve (CN VII), and auditory nerve (CN VIII).

**Fig. S8.**
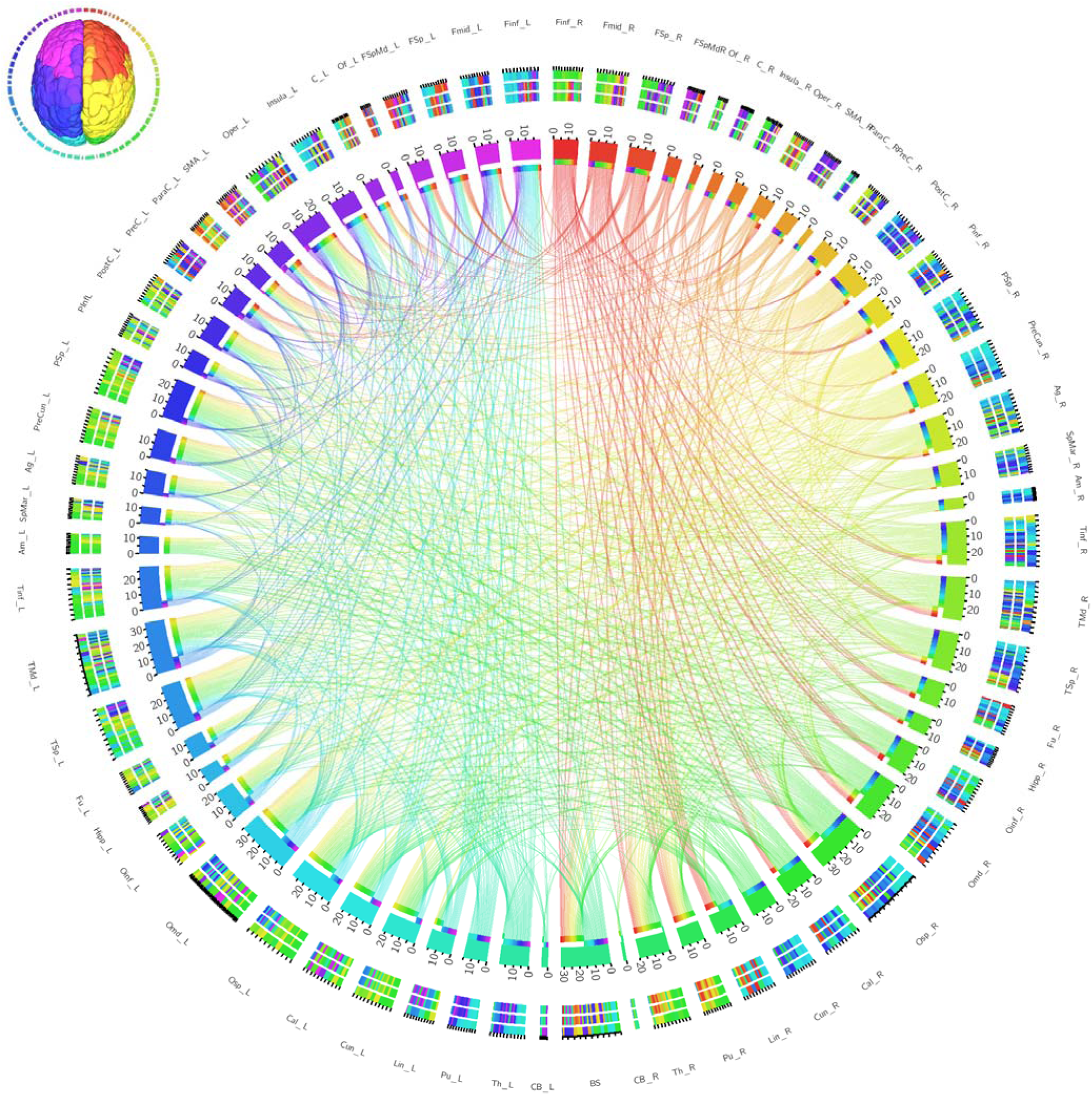
The first level connectogram of the entire human brain connections. The brain is parcellation into regions, and each region is color-coded as shown in the inset figure to the left upper corner. The left side of the connectogram corresponds to the left hemisphere, whereas the right side of the connectogram corresponds to the right hemisphere. The connectogram of the human brain shows a dense network topology between the brain regions, forming a complicated architecture.

**Fig. S9.**
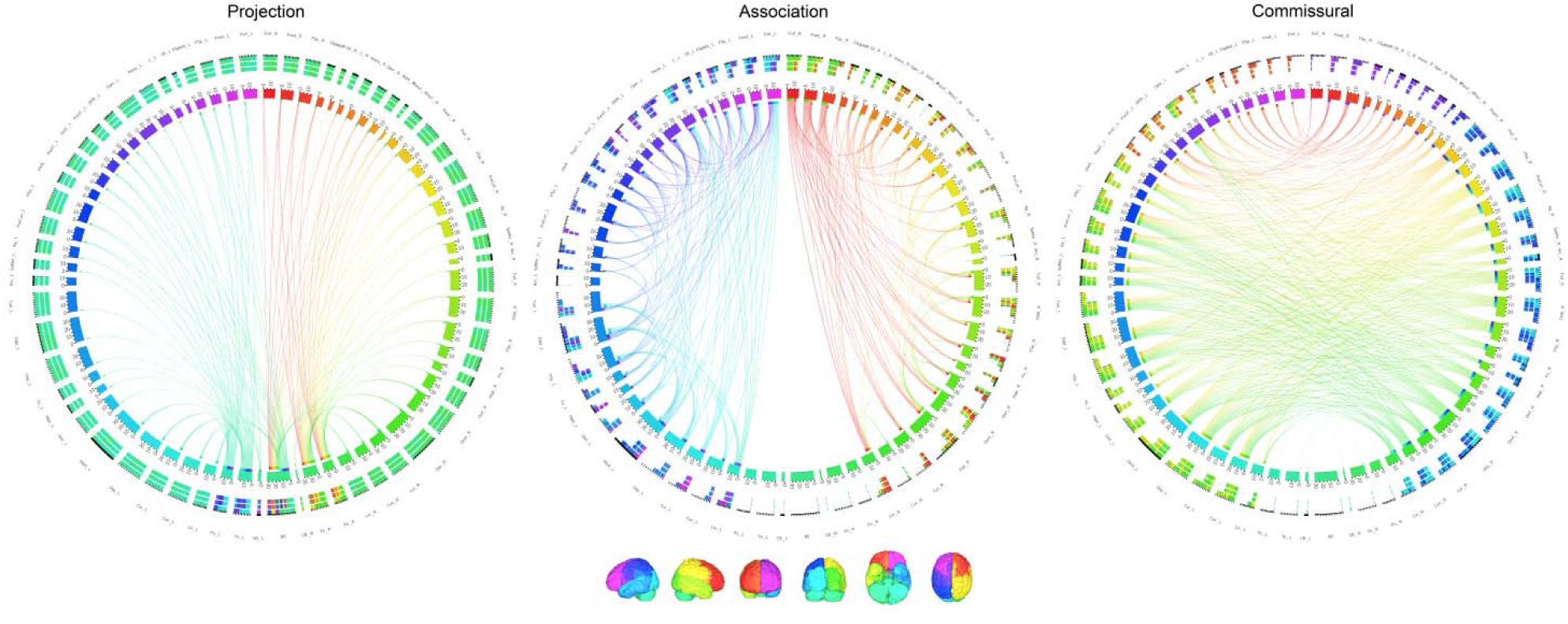
The second level connectograms of the projection, association, and commissural pathways showing the network topology of each pathway system. The projection pathway forms a hub structure in thalamus, putamen, and brainstem. The association pathway forms numerous clusters within each hemisphere. The commissural pathway provides long ranged communication between the two hemispheres.

**Table S1.**
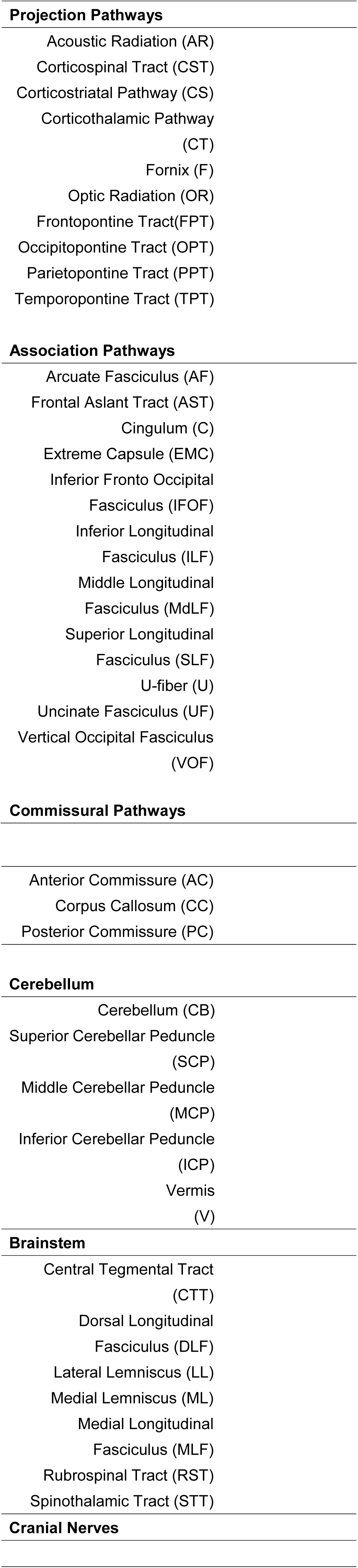
**Abbreviations of the fiber pathways**

**Table S2.**
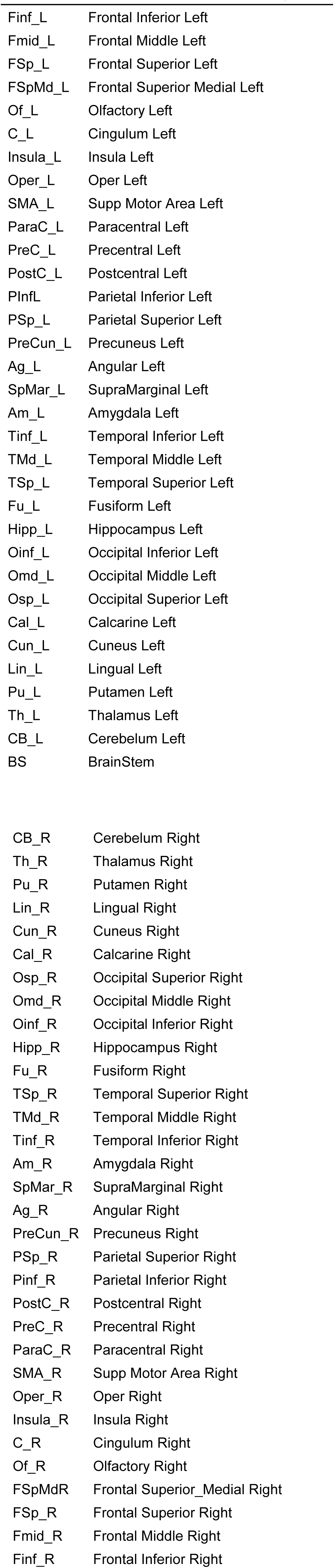
**Abbreviations of the brain regions**

## References

Amunts, K., Lepage, C., Borgeat, L., Mohlberg, H., Dickscheid, T., Rousseau, M.E., Bludau, S., Bazin, P.L., Lewis, L.B., Oros-Peusquens, A.M., Shah, N.J., Lippert, T., Zilles, K., Evans, A.C., 2013. BigBrain: an ultrahigh-resolution 3D human brain model. Science 340, 1472–1475.

Ashburner, J., Friston, K.J., 1999. Nonlinear spatial normalization using basis functions. Hum Brain Mapp 7, 254–266.

Basser, P.J., Pajevic, S., Pierpaoli, C., Duda, J., Aldroubi, A., 2000. In vivo fiber tractography using DT-MRI data. Magn Reson Med 44, 625–632.

Bota, M., Sporns, O., Swanson, L.W., 2015. Architecture of the cerebral cortical association connectome underlying cognition. Proc Natl Acad Sci U S A 112, E2093–2101.

Catani, M., Howard, R.J., Pajevic, S., Jones, D.K., 2002. Virtual in vivo interactive dissection of white matter fasciculi in the human brain. Neuroimage 17, 77–94.

Ding, S.L., Royall, J.J., Sunkin, S.M., Ng, L., Facer, B.A., Lesnar, P., Guillozet-Bongaarts, A., McMurray, B., Szafer, A., Dolbeare, T.A., Stevens, A., Tirrell, L., Benner, T., Caldejon, S., Dalley, R.A., Dee, N., Lau, C., Nyhus, J., Reding, M., Riley, Z.L., Sandman, D., Shen, E., van der Kouwe, A., Varjabedian, A., Write, M., Zollei, L., Dang, C., Knowles, J.A., Koch, C., Phillips, J.W., Sestan, N., Wohnoutka, P., Zielke, H.R., Hohmann, J.G., Jones, A.R., Bernard, A., Hawrylycz, M.J., Hof, P.R., Fischl, B., Lein, E.S., 2016. Comprehensive cellular-resolution atlas of the adult human brain. J Comp Neurol 524, 3127–3481.

Fan, Q., Witzel, T., Nummenmaa, A., Van Dijk, K.R., Van Horn, J.D., Drews, M.K., Somerville, L.H., Sheridan, M.A., Santillana, R.M., Snyder, J., Hedden, T., Shaw, E.E., Hollinshead, M.O., Renvall, V., Zanzonico, R., Keil, B., Cauley, S., Polimeni, J.R., Tisdall, D., Buckner, R.L., Wedeen, V.J., Wald, L.L., Toga, A.W., Rosen, B.R., 2016. MGH-USC Human Connectome Project datasets with ultra-high b-value diffusion MRI. Neuroimage 124, 1108–1114.

Fernandez-Miranda, J.C., Wang, Y., Pathak, S., Stefaneau, L., Verstynen, T., Yeh, F.C., 2015. Asymmetry, connectivity, and segmentation of the arcuate fascicle in the human brain. Brain Struct Funct 220, 1665–1680.

Glasser, M.F., Smith, S.M., Marcus, D.S., Andersson, J.L., Auerbach, E.J., Behrens, T.E., Coalson, T.S., Harms, M.P., Jenkinson, M., Moeller, S., Robinson, E.C., Sotiropoulos, S.N., Xu, J., Yacoub, E., Ugurbil, K., Van Essen, D.C., 2016. The Human Connectome Project’s neuroimaging approach. Nat Neurosci 19, 1175–1187.

Guevara, P., Duclap, D., Poupon, C., Marrakchi-Kacem, L., Fillard, P., Le Bihan, D., Leboyer, M., Houenou, J., Mangin, J.F., 2012. Automatic fiber bundle segmentation in massive tractography datasets using a multi-subject bundle atlas. Neuroimage 61, 1083–1099.

Jeurissen, B., Leemans, A., Tournier, J.D., Jones, D.K., Sijbers, J., 2013. Investigating the prevalence of complex fiber configurations in white matter tissue with diffusion magnetic resonance imaging. Hum Brain Mapp 34, 2747–2766.

Maier-Hein, K., Neher, P., Houde, J.-C., Cote, M.-A., Garyfallidis, E., Zhong, J., Chamberland, M., Yeh, F.-C., Lin, Y.C., Ji, Q., Reddick, W.E., Glass, J.O., Chen, D.Q., Feng, Y., Gao, C., Wu, Y., Ma, J., Renjie, H., Li, Q., Westin, C.-F., Deslauriers-Gauthier, S., Gonzalez, J.O.O., Paquette, M., St-Jean, S., Girard, G., Rheault, F., Sidhu, J., Tax, C.M.W., Guo, F., Mesri, H.Y., David, S., Froeling, M., Heemskerk, A.M., Leemans, A., Bore, A., Pinsard, B., Bedetti, C., Desrosiers, M., Brambati, S., Doyon, J., Sarica, A., Vasta, R., Cerasa, A., Quattrone, A., Yeatman, J., Khan, A.R., Hodges, W., Alexander, S., Romascano, D., Barakovic, M., Auria, A., Esteban, O., Lemkaddem, A., Thiran, J.-P., Cetingul, H.E., Odry, B.L., Mailhe, B., Nadar, M., Pizzagalli, F., Prasad, G., Villalon-Reina, J., Galvis, J., Thompson, P., Requejo, F., Laguna, P., Lacerda, L., Barrett, R., Dell’Acqua, F., Catani, M., Petit, L., Caruyer, E., Daducci, A., Dyrby, T., Holland-Letz, T., Hilgetag, C., Stieltjes, B., Descoteaux, M., 2016. Tractography-based connectomes are dominated by false-positive connections. bioRxiv.

Maier-Hein, K., Neher, P., Houde, J. C., Cote, M. A., Garyfallidis, E., Zhong, J., … & Reddick, W. E., 2016. Tractography-based connectomes are dominated by false-positive connections. bioRxiv 084137.

McNab, J.A., Edlow, B.L., Witzel, T., Huang, S.Y., Bhat, H., Heberlein, K., Feiweier, T., Liu, K., Keil, B., Cohen-Adad, J., Tisdall, M.D., Folkerth, R.D., Kinney, H.C., Wald, L.L., 2013. The Human Connectome Project and beyond: initial applications of 300 mT/m gradients. Neuroimage 80, 234–245.

Meola, A., Comert, A., Yeh, F.C., Sivakanthan, S., Fernandez-Miranda, J.C., 2016a. The nondecussating pathway of the dentatorubrothalamic tract in humans: human connectome-based tractographic study and microdissection validation. J Neurosurg 124, 1406–1412.

Meola, A., Comert, A., Yeh, F.C., Stefaneanu, L., Fernandez-Miranda, J.C., 2015. The controversial existence of the human superior fronto-occipital fasciculus: Connectome-based tractographic study with microdissection validation. Hum Brain Mapp 36, 4964–4971.

Meola, A., Yeh, F.C., Fellows-Mayle, W., Weed, J., Fernandez-Miranda, J.C., 2016b. Human Connectome-Based Tractographic Atlas of the Brainstem Connections and Surgical Approaches. Neurosurgery 79, 437–455.

Mori, S., Crain, B.J., Chacko, V.P., van Zijl, P.C., 1999. Three-dimensional tracking of axonal projections in the brain by magnetic resonance imaging. Ann Neurol 45, 265–269.

Mori, S., Oishi, K., Faria, A.V., 2009. White matter atlases based on diffusion tensor imaging. Curr Opin Neurol 22, 362–369.

Mori, S., Oishi, K., Jiang, H., Jiang, L., Li, X., Akhter, K., Hua, K., Faria, A.V., Mahmood, A., Woods, R., Toga, A.W., Pike, G.B., Neto, P.R., Evans, A., Zhang, J., Huang, H., Miller, M.I., van Zijl, P., Mazziotta, J., 2008. Stereotaxic white matter atlas based on diffusion tensor imaging in an ICBM template. Neuroimage 40, 570–582.

Peng, H., Orlichenko, A., Dawe, R.J., Agam, G., Zhang, S., Arfanakis, K., 2009. Development of a human brain diffusion tensor template. Neuroimage.

Reveley, C., Seth, A.K., Pierpaoli, C., Silva, A.C., Yu, D., Saunders, R.C., Leopold, D.A., Ye, F.Q., 2015. Superficial white matter fiber systems impede detection of long-range cortical connections in diffusion MR tractography. Proc Natl Acad Sci U S A 112, E2820–2828.

Setsompop, K., Kimmlingen, R., Eberlein, E., Witzel, T., Cohen-Adad, J., McNab, J.A., Keil, B., Tisdall, M.D., Hoecht, P., Dietz, P., Cauley, S.F., Tountcheva, V., Matschl, V., Lenz, V.H., Heberlein, K., Potthast, A., Thein, H., Van Horn, J., Toga, A., Schmitt, F., Lehne, D., Rosen, B.R., Wedeen, V., Wald, L.L., 2013. Pushing the limits of in vivo diffusion MRI for the Human Connectome Project. Neuroimage 80, 220–233.

Sporns, O., 2014. Contributions and challenges for network models in cognitive neuroscience. Nat Neurosci 17, 652–660.

Thiebaut de Schotten, M., Ffytche, D.H., Bizzi, A., Dell’Acqua, F., Allin, M., Walshe, M., Murray, R., Williams, S.C., Murphy, D.G., Catani, M., 2011. Atlasing location, asymmetry and inter-subject variability of white matter tracts in the human brain with MR diffusion tractography. Neuroimage 54, 49–59.

Thomas, C., Ye, F.Q., Irfanoglu, M.O., Modi, P., Saleem, K.S., Leopold, D.A., Pierpaoli, C., 2014. Anatomical accuracy of brain connections derived from diffusion MRI tractography is inherently limited. Proc Natl Acad Sci U S A 111, 16574–16579.

Van Essen, D.C., Smith, S.M., Barch, D.M., Behrens, T.E., Yacoub, E., Ugurbil, K., Consortium, W.U.-M.H., 2013. The WU-Minn Human Connectome Project: an overview. Neuroimage 80, 62–79.

Van Essen, D.C., Ugurbil, K., Auerbach, E., Barch, D., Behrens, T.E., Bucholz, R., Chang, A., Chen, L., Corbetta, M., Curtiss, S.W., Della Penna, S., Feinberg, D., Glasser, M.F., Harel, N., Heath, A.C., Larson-Prior, L., Marcus, D., Michalareas, G., Moeller, S., Oostenveld, R., Petersen, S.E., Prior, F., Schlaggar, B.L., Smith, S.M., Snyder, A.Z., Xu, J., Yacoub, E., 2012. The Human Connectome Project: a data acquisition perspective. Neuroimage 62, 2222–2231.

Wang, X., Pathak, S., Stefaneanu, L., Yeh, F.C., Li, S., Fernandez-Miranda, J.C., 2016. Subcomponents and connectivity of the superior longitudinal fasciculus in the human brain. Brain Struct Funct 221, 2075–2092.

Wang, Y., Fernandez-Miranda, J.C., Verstynen, T., Pathak, S., Schneider, W., Yeh, F.C., 2013. Rethinking the role of the middle longitudinal fascicle in language and auditory pathways. Cereb Cortex 23, 2347–2356.

Wedeen, V.J., Rosene, D.L., Wang, R., Dai, G., Mortazavi, F., Hagmann, P., Kaas, J.H., Tseng, W.Y., 2012. The geometric structure of the brain fiber pathways. Science 335, 1628–1634.

Yeh, F.C., Tseng, W.Y., 2011. NTU-90: a high angular resolution brain atlas constructed by q-space diffeomorphic reconstruction. Neuroimage 58, 91–99.

Yeh, F.C., Verstynen, T.D., Wang, Y., Fernandez-Miranda, J.C., Tseng, W.Y., 2013. Deterministic diffusion fiber tracking improved by quantitative anisotropy. PLoS ONE 8, e80713.

Yeh, F.C., Wedeen, V.J., Tseng, W.Y., 2010. Generalized q-sampling imaging. IEEE Trans Med Imaging 29, 1626–1635.

Yoshino, M., Abhinav, K., Yeh, F.C., Panesar, S., Fernandes, D., Pathak, S., Gardner, P.A., Fernandez-Miranda, J.C., 2016. Visualization of Cranial Nerves Using High-Definition Fiber Tractography. Neurosurgery 79, 146–165.

Zhang, S., Peng, H., Dawe, R.J., Arfanakis, K., 2011. Enhanced ICBM diffusion tensor template of the human brain. Neuroimage 54, 974–984.

Zhang, W., Olivi, A., Hertig, S.J., van Zijl, P., Mori, S., 2008. Automated fiber tracking of human brain white matter using diffusion tensor imaging. Neuroimage 42, 771–777.

